# A pseudovirus system enables deep mutational scanning of the full SARS-CoV-2 spike

**DOI:** 10.1101/2022.10.13.512056

**Authors:** Bernadeta Dadonaite, Katharine H D Crawford, Caelan E Radford, Ariana G Farrell, Timothy C Yu, William W Hannon, Panpan Zhou, Raiees Andrabi, Dennis R Burton, Lihong Liu, David D. Ho, Richard A. Neher, Jesse D Bloom

## Abstract

A major challenge in understanding SARS-CoV-2 evolution is interpreting the antigenic and functional effects of emerging mutations in the viral spike protein. Here we describe a new deep mutational scanning platform based on non-replicative pseudotyped lentiviruses that directly quantifies how large numbers of spike mutations impact antibody neutralization and pseudovirus infection. We demonstrate this new platform by making libraries of the Omicron BA.1 and Delta spikes. These libraries each contain ~7000 distinct amino-acid mutations in the context of up to ~135,000 unique mutation combinations. We use these libraries to map escape mutations from neutralizing antibodies targeting the receptor binding domain, N-terminal domain, and S2 subunit of spike. Overall, this work establishes a high-throughput and safe approach to measure how ~10^5^ combinations of mutations affect antibody neutralization and spike-mediated infection. Notably, the platform described here can be extended to the entry proteins of many other viruses.

## Introduction

The spike protein is the key target of neutralizing antibodies against SARS-CoV-2. Unfortunately, spike has undergone rapid evolution which has eroded the potency of serum neutralization and enabled escape from most monoclonal antibodies (Cao et al., 2022; Liu et al., 2022; Wang et al., 2022b, 2022a). Deep mutational scanning experiments can prospectively measure the effects of large numbers of mutations even before they emerge in viral variants, and therefore have been a valuable tool for rapidly interpreting how newly observed mutations in the spike affect antibody binding and protein folding or function (Cao et al., 2022; Starr et al., 2021b, 2021a). The high-throughput nature of deep mutational scanning experiments has also enabled the generation of huge datasets that can inform computational methods for predicting the antigenic properties of possible future viral variants (Cao et al., 2022; Greaney et al., 2022).

However, prior deep mutational scanning of the SARS-CoV-2 spike has been limited to either solely focusing on the receptor-binding domain (RBD) (Cao et al., 2022; Greaney et al., 2021; Starr et al., 2020), other subdomains (Ouyang et al., 2022; Tan et al., 2022) or just a small number of mutations across spike (Javanmardi et al., 2021). Furthermore, all previous spike deep mutational scanning experiments have been based on cell-surface display using either yeast (Starr et al., 2022, 2020) or mammalian cells (Javanmardi et al., 2021; Ouyang et al., 2022; Tan et al., 2022), and therefore are limited to measuring antibody binding rather than neutralization, despite the fact that neutralization is thought to be a more relevant correlate of protection (Feng et al., 2021; Gilbert et al., 2022).

Here we describe a new deep mutational scanning platform that directly measures how mutations affect cellular infection and antibody neutralization in the context of the full SARS-CoV-2 spike pseudotyped on non-replicative lentiviral particles. The key innovation behind the platform is a two-step pseudovirus generation protocol that enables creation of large pseudovirus libraries with a link between the lentiviral genotype and the particular spike protein variant on the pseudovirus’s surface. We demonstrate that this new platform can be used to create large genotype-phenotype linked pseudovirus libraries and map how mutations to spike affect both cellular infection and neutralization by antibodies targeting diverse regions of spike, including the RBD, N-terminal domain (NTD), and S2 subunit.

## Results

### Producing pseudoviruses with genotype-phenotype link

To characterize thousands of mutations in spike glycoprotein, we first established a lentiviral pseudotyping platform that maintains a genotype-phenotype link between the lentiviral genome and the spike variant on the virion’s surface. Lentiviral spike-pseudotyping usually involves transfection of a backbone that carries a reporter gene flanked by the lentiviral long terminal repeats (LTRs), helper plasmids that code for structural and nonstructural genes required for the lentiviral life cycle, and an expression plasmid that codes for the spike variant of interest (Crawford et al., 2020; Cronin et al., 2005; Naldini et al., 1996). When these components are transfected into producer cells, virions are formed that carry lentiviral genomes and display spikes on their surface. However, because genome incorporation into a virion does not depend on the expressed spike, there is no link between the virion’s genotype and the phenotype of the spike on its surface. The absence of a genotypephenotype link is not problematic when only a single spike variant is used for transfection — however, it precludes deep mutational scanning studies that involve studying thousands of variants in a single pooled experiment.

To create a lentiviral genotype-phenotype link, we first generated a lentivirus backbone with the following key elements (Figure 1A): (1) we restored the ability of the lentivirus to transcribe its full genome after integration by repairing the 3’ LTR deletion present in traditional lentivirus vectors (OhAinle et al., 2018; Zufferey et al., 1998), (2) we placed spike in the lentivirus backbone under an inducible promoter, (3) we added a second constitutive promoter driving both a fluorescent reporter (ZsGreen) and a puromycin resistance gene.

**Figure 1.**
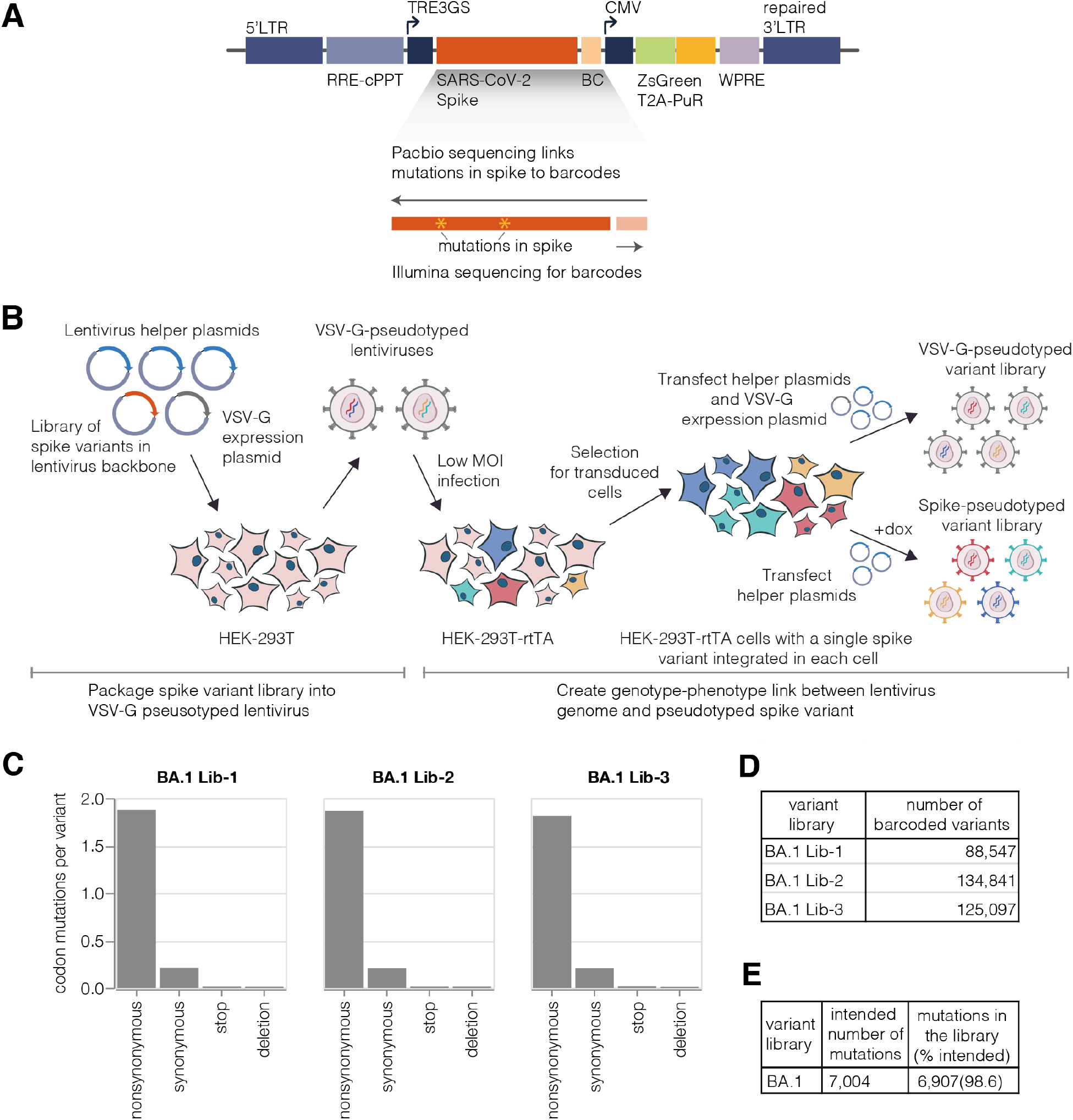
Deep mutational scanning platform for spike. **(A)** Lentivirus backbone used for deep mutational scanning. The backbone contains functional lentiviral 5’ and 3’ long terminal repeat (LTR) regions. The spike gene is under an inducible tet response element 3rd generation (TRE3G) promoter, and there is a 16 nucleotide barcode (BC) downstream of the stop codon. A CMV promoter drives expression of reporter ZsGreen gene that is linked to a puromycin resistance gene (PuR) via a T2A linker. The backbone also contains a woodchuck hepatitis virus posttranscriptional regulatory element (WPRE), Rev response element (RRE), and a central polypurine tract (cPPT). **(B)** Approach for creating genotype-phenotype linked lentivirus libraries. HEK-293T cells are transfected with spike-carrying lentivirus backbone, VSV-G expression plasmid and lentiviral helper plasmids to generate VSV-G-pseudotyped lentiviruses. These viruses are used to transduce reverse tetracycline-controlled transactivator (rtTA) expressing HEK-293T cells at low multiplicity of infection (MOI), and successfully transduced cells are selected using puromycin. Selected cells can be transfected with helper plasmids and a VSV-G expression plasmid to produce VSV-G-pseudotyped viruses carrying all genomes present in the deep mutational library or selected cells can be induced with doxycycline (dox) to express spike and transfected with only the helper plasmids to generate spike-pseudotyped lentiviruses that have a genotype-phenotype link. **(C)** Average number of mutations per barcoded spike in BA.1 libraries. **(D)** Total number of barcoded variants in each BA.1 library. **(E)** The coverage of intended mutations across all BA.1 libraries.

Next, we developed a multi-step protocol that creates a genotype-phenotype link by ensuring that each producer cell only expresses a single variant of spike (Figure 1B). In the first step of this protocol, we transfect cells with the spike-encoding backbone, a VSV-G expression plasmid, and the necessary helper plasmids. This produces non-genotype-phenotype-linked VSV-G-pseudotyped lentiviruses that we use to infect target cells at low multiplicity of infection, so that most infected cells receive no more than one lentiviral genome. Next we select for cells with integrated lentiviral genomes using puromycin, which yields a population of cells where each cell stores only a single spike variant. The spike is under an inducible promoter, which is only activated by addition of doxycycline. To produce virions, we induce spike expression with doxycycline and transfect the helper plasmids necessary to produce lentiviruses. We validated that this approach can be used to generate genotype-phenotype linked spike-pseudotyped viruses with titers >10^5^ transduction units per ml (Figure S1A). We can further increase viral titers by ~5-10 fold by infecting cells in the presence of a putative IFITM3 inhibitor amphotericin B (Lin et al., 2013), as has been reported previously (Peacock et al., 2021; Zhao et al., 2020; Zheng et al., 2020) (Figure S1B).

### Design of mutations in SARS-CoV-2 spike deep mutational scanning library

Rather than create deep mutational scanning libraries containing all possible amino-acid mutants of spike, we chose to introduce only mutations that seem likely to arise during natural evolution and yield a functional spike protein. We had two rationales for designing our libraries in this way: (1) it reduces the total number of mutations that need to be included in the library, and (2) it increases the probability that variants with multiple mutations will remain functional by reducing the fraction of mutations that are highly deleterious.

Specifically, we included only mutations that have been observed in spike sequences deposited on the GISAID database (Khare et al., 2021), reasoning that these mutations would represent mostly functional spike proteins. We introduced mutations at a higher frequency when they have emerged in spike independently many times according to the pre-built SARS-CoV-2 phylogenies from UShER (Turakhia et al., 2021). Finally, we included every possible amino acid change at sites in spike that are evolving under positive selection (Maher et al., 2022). We also included deletions at sites where such mutations are observed frequently in natural SARS-CoV-2 evolution. In total, our library design targeted 7,004 mutations in the BA.1 spike and 6,852 mutations in the Delta spike.

To introduce these mutations in the spike gene we used a PCR-based mutagenesis method with a primer pool containing the desired mutations (Bloom, 2014). Importantly, this method introduces multiple mutations in each spike variant: we targeted ~2 to 3 codon mutations per variant, ensuring the effects of most mutations are measured in multiple genetic backgrounds. The mutated spike genes were then barcoded with 16 random nucleotides placed downstream of the spike-coding sequence (Figure 1A), and cloned into the lentivirus backbone. As described below, after integration of the libraries into cells, these barcodes can be linked to the full set of mutations in each spike variant to facilitate downstream sequencing (Hiatt et al., 2010; Matreyek et al., 2018).

### Production of pseudotyped BA.1 and Delta spike deep mutational scanning libraries

We used the genotype-phenotype linked pseudovirus production strategy in Figure 1B to make BA.1 and Delta deep mutational scanning libraries. We created three independent BA.1 libraries each containing ~100,000 barcoded variants and two independent Delta libraries each containing ~50,000 barcoded variants (Figures 1E, S2A; a “barcoded variant” is a spike with a unique nucleotide barcode and some random mutation set; different barcoded variants usually but not always contain different mutations). After integrating the libraries into cells at low multiplicity of infection, we generated VSV-G-pseudotyped lentivirus from these cells by co-transfecting a plasmid expressing VSV-G alongside the other lentiviral helper plasmids (Figure 1B, top right). The use of VSV-G-pseudotyped virus ensures that we generate infectious lentiviral virions from all integrated backbones regardless of whether they encode a functional spike mutant. We then infected this VSV-G-pseudotyped lentivirus into a new round of cells, and performed long-read PacBio sequencing to link the barcodes to the full set of spike mutations for each variant. We performed the PacBio barcode-mutation linking after integration into cells because recombination of the pseudodiploid lentiviral genome during integration (Jetzt et al., 2000; Schlub et al., 2010) means the barcode-mutation pairings may be different in the integrated cells to those in the original lentiviral backbone plasmids (Hill et al., 2018). Importantly, linking barcodes to spike variants allows us to use short-read Illumina sequencing of the barcode to obtain the full spike genotype in all subsequent experiments.

Overall, the sequencing revealed that we had successfully introduced ~99% of the targeted mutations in the BA.1 and Delta spike libraries (Figures 1E, S2B). The barcoded variants in the BA.1 libraries had on average ~2 codon mutations per spike, while the variants in the Delta libraries had ~3 codon mutations per spike (Figures 1C, S2C). The number of mutations per variant is roughly Poisson distributed, so some variants had zero or one mutation, while others had many more (Figure S3).

We then generated the actual spike-pseudotyped deep mutational scanning libraries from the variants stored at single copy in the cells (Figure 1B, lower right). We calculated a functional score for each variant based on its relative frequency in the spike-versus VSV-G-pseudotyped libraries. Positive functional scores indicate spike variants mediate pseudovirus infection better than the parental spike, whereas negative functional scores indicate worse pseudovirus infection. As expected, spike variants with premature stop codons had highly negative functional scores, while unmutated and synonymously mutated spike variants had functional scores close to zero (Figures 2, S2D). Some variants with nonsynonymous mutations had functional scores close to zero, while others had more negative scores, reflecting the fact that some but not all nonsynonymous mutations are deleterious (Figures 2, S2D; recall that our library design protocol preferentially introduced nonsynonymous mutations expected to yield functional spikes). Variants with multiple nonsynonymous mutations tended to have lower functional scores than variants with just one nonsynonymous mutation (Figures 2, S2D), reflecting the cost of accumulating multiple often mildly deleterious mutations.

**Figure 2.**
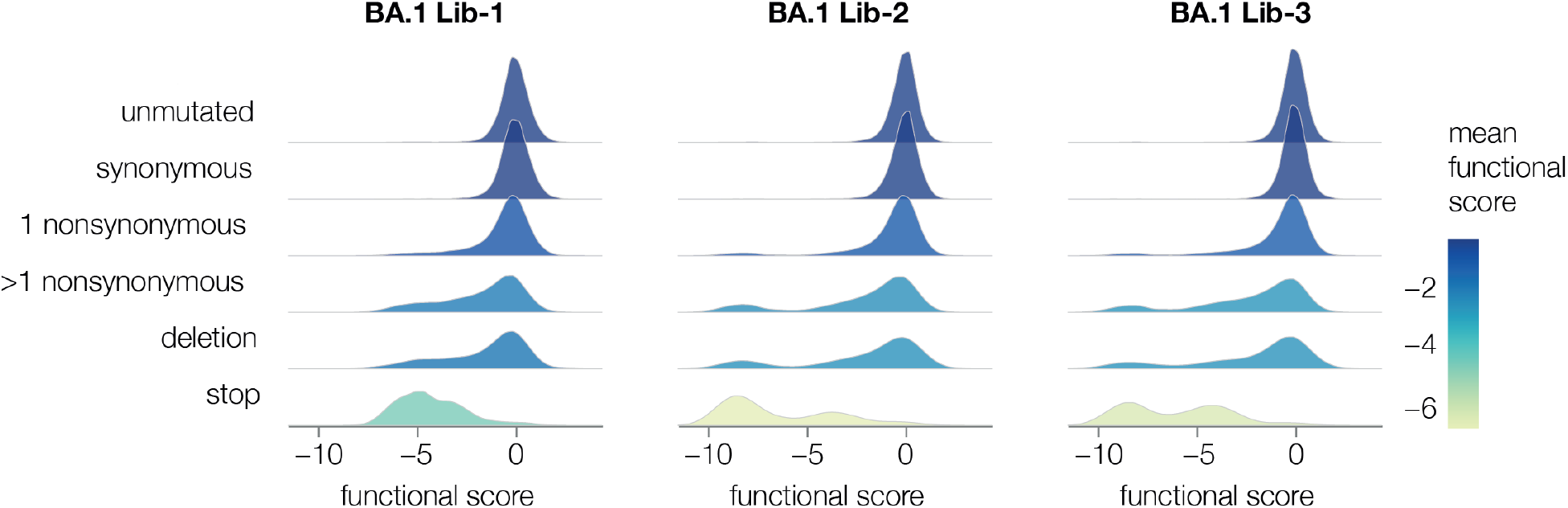
Some types of mutations tend to impair spike-mediated pseudovirus infection. For each barcoded spike variant, we compute a functional score that reflects how well that spike mediates pseudovirus infection relative to the unmutated spike: negative scores indicate impaired infection, positive scores indicate improved infection. The plots show the distribution of functional scores across all variants in each of the three BA.1 libraries for different categories of variants, with each distribution colored by the mean functional score for that variant type.

### Use of an absolute standard to measure viral neutralization by deep sequencing

Traditional neutralization assays measure the infectivity of a single virus variant at multiple antibody concentrations. Deep sequencing can measure the relative infectivities of many viral variants in pooled infections in the presence of an antibody. However, to convert the relative infectivities measured by deep sequencing into actual neutralization values, it is necessary to have an absolute standard that does not vary in its infectivity as a function of antibody concentration (Figure 3A). To enable such measurements in our experiments, we added a barcoded VSV-G-pseudotyped virus into our libraries. Importantly, this VSV-G-pseudotyped virus is not neutralized by any of the spike-binding antibodies (Figures 3A, S4A), and so the counts for VSV-G barcodes provide an absolute neutralization standard. To calculate the non-neutralized fraction for each viral variant, we simply compute the change in its barcode frequencies relative to the VSV-G standard (Figure 3B).

**Figure 3.**
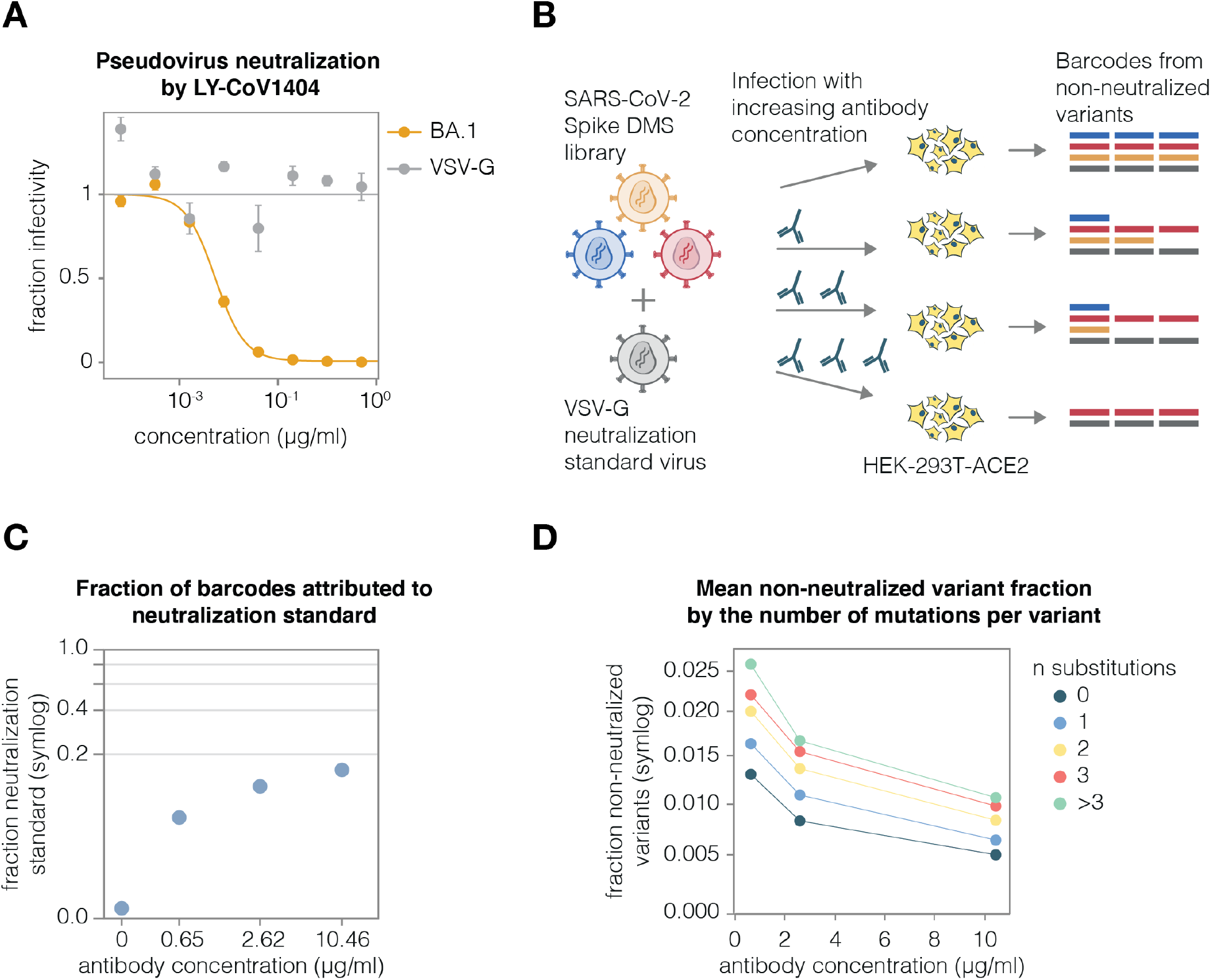
A VSV-G standard enables measurement of absolute neutralization by deep sequencing. **(A)** Neutralization assay demonstrating that BA.1-spike-pseudotyped lentivirus is neutralized by antibody LY-CoV1404, but the VSV-G-pseudotyped neutralization standard is not. **(B)** Use of the VSV-G standard to measure absolute neutralization. Deep mutational scanning libraries are mixed with VSV-G neutralization standard. The virus mixture is incubated with a no-antibody control or increasing antibody concentrations and infected into ACE2-expressing 293T cells. After ~12 hours viral genomes are recovered, barcodes are sequenced, and absolute neutralization of each variant is computed by comparing its barcode counts to those from the VSV-G standard. **(C)** Fraction of barcodes derived from the VSV-G neutralization standard in infections with increasing LY-CoV1404 concentrations. **(D)** BA.1 deep mutational scanning library non-neutralized fractions averaged across variants with different numbers of amino-acid mutations at differentLY-CoV1404 concentrations.Note panels C and D use a symlog scale.

To validate this approach, we added the VSV-G absolute standard at ~1% of our BA.1 library titers and incubated the virus library with increasing concentrations of the LY-CoV1404 antibody, as schematized in Figure 3B. We then infected the library into ACE2-expressing target cells overnight, recovered viral genomes, and quantified the abundance of each viral barcode using deep sequencing. As expected, the fraction of VSV-G standard reads increased with antibody concentration because fewer spike variants could still infect in the presence of antibody (Figure 3C). We then calculated the non-neutralized fraction for each viral variant in our libraries after selection at different concentrations of the antibody. As expected, increasing antibody concentrations led to decreased non-neutralized fraction averaged over variants (Figure 3D). Notably, variants with greater number of substitutions had higher non-neutralized fractions, as expected if some substitutions escape the antibody.

### Mapping antibody escape using a full spike deep mutational scanning system

To demonstrate that pseudovirus-based deep mutational scanning can map escape from neutralizing antibodies targeting any region of spike, we chose a set of BA.1-neutralizing antibodies that bind distinct regions of spike: RBD-binding LY-CoV1404, NTD-binding 5-7, and S2-binding CC67.105 (Cerutti et al., 2021; Westendorf et al., 2022; Zhou et al., 2022). Note that LY-CoV1404, also known as bebtelovimab, is one of the few clinically approved antibodies that retains potency against BA.1, BA.2 and other major Omicron lineages (Wang et al., 2022a; Westendorf et al., 2022).

We first mapped escape from LY-CoV1404, applying the approach in Figure 3B to our three independent BA.1 libraries, and performing a technical replicate for one library. We used a biophysical model to decompose the measurements for spike variants in our libraries (some of which are multiply mutated) into escape scores for individual mutations (Yu et al., 2022). These mutation escape scores correlated well among technical and biological replicates (Figure 4A). As expected, the key LY-CoV1404 escape sites were in the antibody’s previously described epitope in the RBD (Westendorf et al., 2022), which spans sites 439-452 and 498-501 (Figure 4B-D and https://dms-vep.github.io/SARS-CoV-2_Omicron_BA.1_spike_DMS_mAbs/LyCoV-1404_escape_plot.html).

**Figure 4.**
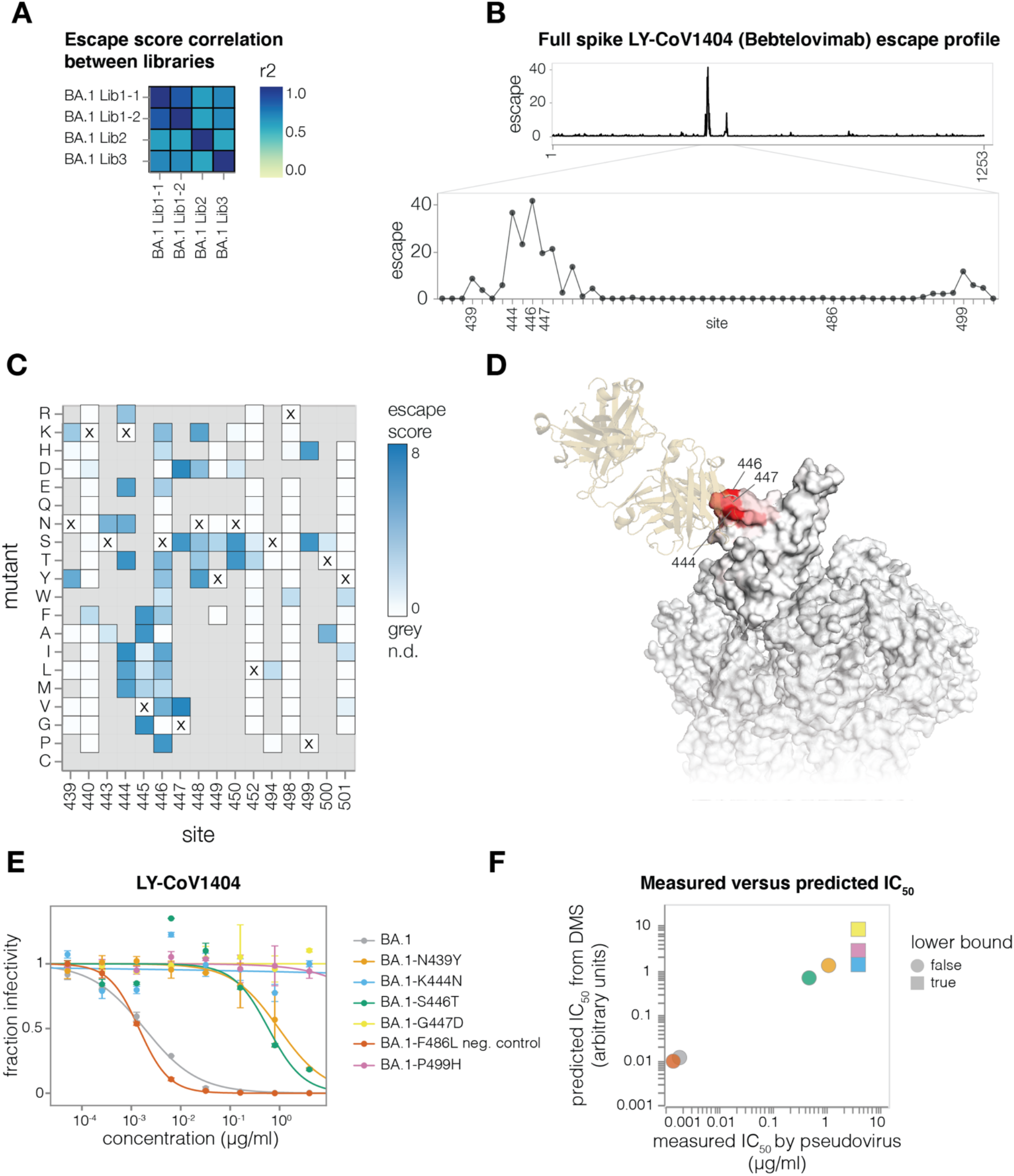
Antibody LY-CoV1404 escape mapping. **(A)** Correlation of mutation escape scores between technical replicates (BA1 Lib-1.1 and BA1 Lib-1.2) and biological replicates (BA1 Lib-1, BA1 Lib-2, BA1 Lib-3). **(B)** Total escape scores at each site in the BA.1 spike, and a zoomed-in plot showing the key escape sites. Sites of mutations chosen for validation experiments are labeled on the x-axis. **(C)** Heatmap of mutation escape scores at key sites. Residues marked with X are the wild-type amino acids in BA.1. Amino acids not present in our libraries are shown in gray. An interactive heatmap for the entirety of spike is at https://dms-vep.github.io/SARS-CoV-2_Omicron_BA.1_spike_DMS_mAbs/LyCoV-1404_escape_plot.html **(D)** Surface representation of spike coloured by sum of escape scores at that site. LY-CoV1404 antibody is in yellow. Only the antibody-bound protomer is coloured. PDB IDs 7MMO and 6XM4 were aligned to make this structure. **(E)** Validation pseudovirus neutralization assays of the indicated BA.1 spike mutants against the LY-CoV1404 antibody. **(F)** Correlation between predicted IC50 values from deep mutational scanning (DMS) data versus the IC50 values measured in the validation assays in panel (E). The points are colored as in panel (E). Lower bound indicates that the antibody did not neutralize at the highest concentration tested in the validation neutralization assay. Site numbering in all plots is based on the Wuhan-Hu-1 sequence.

However, our deep mutational scanning emphasizes that only some mutations at these sites escape LY-CoV1404 neutralization. For instance, many amino acid mutations at site 446 strongly escape LY-CoV1404, but mutating this site from G (the identity in Wuhan-Hu-1) to S (the identity in BA.1 and BA.2.75) does not have a large effect. This observation emphasizes the somewhat serendipitous nature of the preserved potency of LY-CoV1404. However, this antibody may soon be escaped because sub-variants of BA.5 and BA.2.75 with mutations in the key escape site of K444 are increasingly being detected (Cao et al., 2022; Chen et al., 2022).

To validate the LY-CoV1404 deep mutational scanning, we cloned a set of mutations in the BA.1 spike with a range of effects in the deep mutational scanning data and performed standard pseudovirus neutralization assays (Figure 4E). All the tested mutations exhibited neutralization phenotypes consistent with those measured in the deep mutational scanning. Furthermore, the neutralization assay IC50 values correlated well with those predicted by our biophysical model (Yu et al., 2022) parameterized by the deep mutational scanning data (Figure 4F).

We also compared the full spike LY-CoV1404 deep mutational scanning measurements to results from our previously described yeast-display system for deep mutational scanning of only the RBD (Starr et al., 2022, 2020). The escape scores between the two experimental approaches correlated well (Figure S5A) and both methods identified the same epitope (Figure S5B).

To show that we can map escape from non-RBD-targeting antibodies, we next mapped the NTD-targeting 5-7 antibody (Cerutti et al., 2021). This antibody targets an epitope outside the defined antigenic supersite in NTD and is one of the few NTD-targeting antibodies isolated pre-Omicron that still retains some potency against Omicron variants (Cerutti et al., 2021; Liu et al., 2022; McCallum et al., 2021). The deep mutational scanning showed that the key escape sites for 5-7 were in a hydrophobic pocket next to the N4 loop (site 172-178) (Figure 5A-C and https://dms-vep.github.io/SARS-CoV-2_Omicron_BA.1_spike_DMS_mAbs/NTD_5-7_escape_plot.html), consistent with prior structural characterization of this antibody’s epitope (Cerutti et al., 2021). In addition, deletions in 167-171 ß-sheet, as well as mutations at the base of the adjacent loops such as G103 and V126 also escaped antibody 5-7 (Figure 5A-B). We validated these deep mutational scanning results by performing individual neutralization assays with pseudoviruses containing L176K, S172N and G103F mutations (Figure 5D), all of which had the expected effect of completely escaping neutralization (Figure 5E).

**Figure 5.**
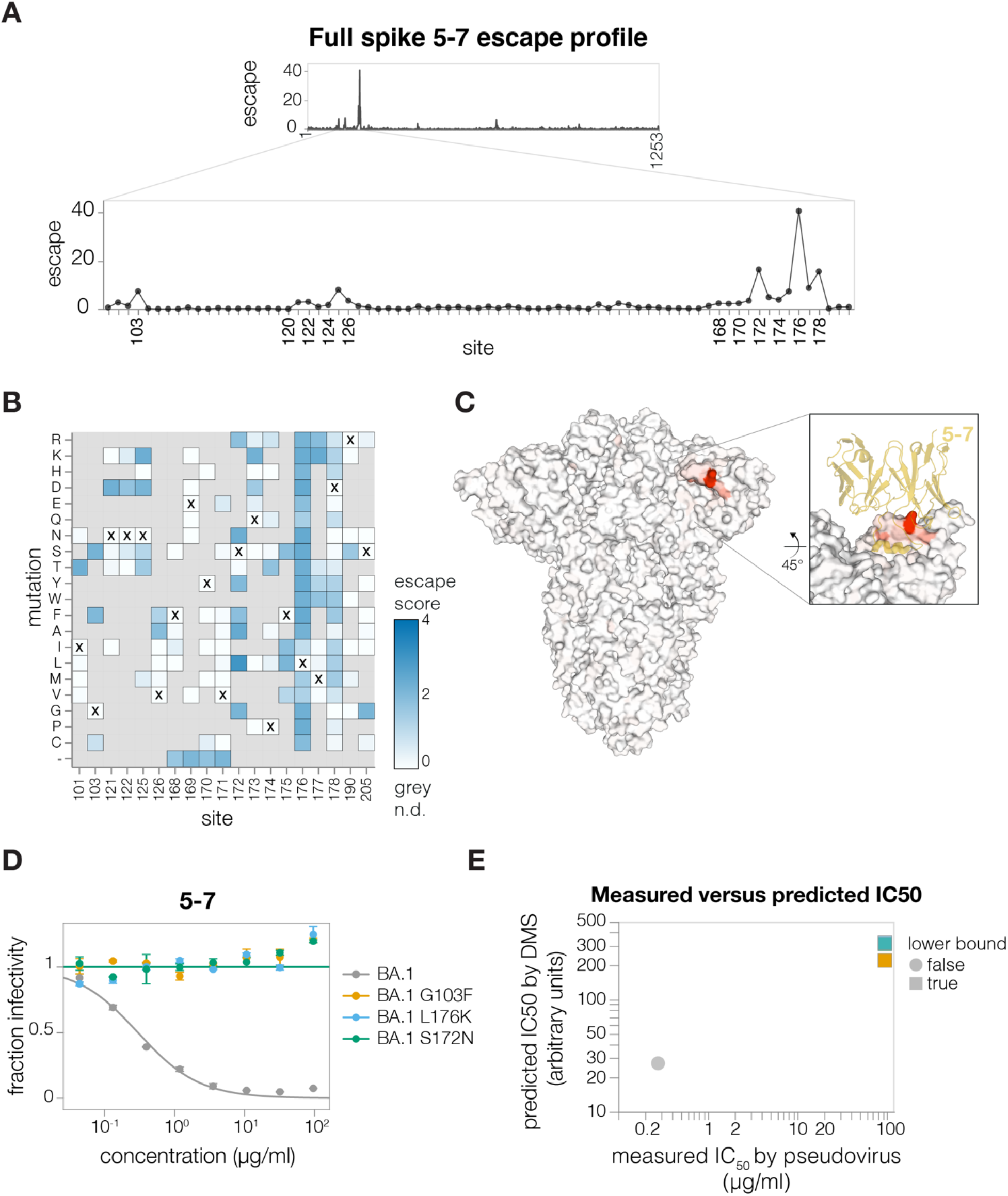
Antibody 5-7 escape mapping. **(A)** Total escape scores for each site in the BA.1 spike and a zoomed-in plot showing the key escape sites. **(B)** Heatmap of mutation escape scores at key sites. Residues marked with X are the wild-type amino acids in BA.1. Amino acids not present in our libraries are shown in gray. An interactive version of this plot for the entirety of spike is at https://dms-vep.github.io/SARS-CoV-2_Omicron_BA.1_spike_DMS_mAbs/NTD_5-7_escape_plot.html **(C)** Surface representation of spike coloured by the sum of escape scores at that site. Antibody 5-7 is shown in yellow in the inset. PDB ID: 7RW2. **(D)** Validation pseudovirus neutralization assays of the indicated BA.1 spike mutants against antibody 5-7. **(E)** Correlation between predicted IC50 values from deep mutational scanning (DMS) data versus the IC50 values measured in the validation assays in (F). Lower bound indicates that the antibody did not neutralize at the highest concentration tested in the validation neutralization assay. Site numbering in all plots is based on the Wuhan-Hu-1 sequence.

We next applied the full spike deep mutational scanning to S2 stem-helix targeting antibodies CC9.104 and CC67.105, which were isolated using both the SARS-CoV-2 and Middle East Respiratory Syndrome coronavirus (MERS-CoV) spike proteins as baits (Zhou et al., 2022). Both CC9.104 and CC67.105 broadly neutralize SARS-related coronaviruses, and CC9.104 also retains some potency against MERS-CoV. As expected, our deep mutational scanning showed that escape sites for both antibodies cluster in the S2 stem-helix region (Figure 6A-E; https://dms-vep.github.io/SARS-CoV-2_Omicron_BA.1_spike_DMS_mAbs/CC67.105_escape_plot.html and https://dms-vep.github.io/SARS-CoV-2_Omicron_BA.1_spike_DMS_mAbs/CC9.104_escape_plot.html). Our data also explain why only CC9.104 neutralizes MERS-CoV. The deep mutational scanning shows that the CC67.105 epitope centers on sites D1146, D1153 and F1156 (Figures 6B, 6D), and consistent with the deep mutational scanning, mutating these sites leads to complete escape in validation neutralization assays (Figure 6G). By contrast, the deep mutational scanning shows that while CC9.104’s epitope also includes sites D1153 and F1156, mutations at site D1146 cause only modest or no escape (Figures 6C, 6E), and validation neutralization assays again confirm these deep mutational scanning results (Figures 6G). Notably, sites D1153 and F1156 are conserved between SARS-CoV-2 and MERS-CoV S2 stem-helix regions, but site D1146 is mutated to isoleucine in MERS-CoV (Figure 6F). Based on our deep mutational scanning data we expect changes at site D1146 to have significant effects on escape from the CC67.105 antibody but only modest effects on escape from the CC9.104 antibody. While our deep mutational scanning libraries do not include isoleucine at site D1146 (which is found in MERS-CoV), we confirmed that other mutations at D1146 lead to complete escape from CC67.105 and do not substantially impact neutralization by CC9.104 (Figure 6G). Note that site D1163 is also mutated to isoleucine in MERS-CoV and both antibodies show some escape at that site, which may explain why CC9.104’s potency against MERS-CoV is lower than against SARS-CoV-2.

**Figure 6.**
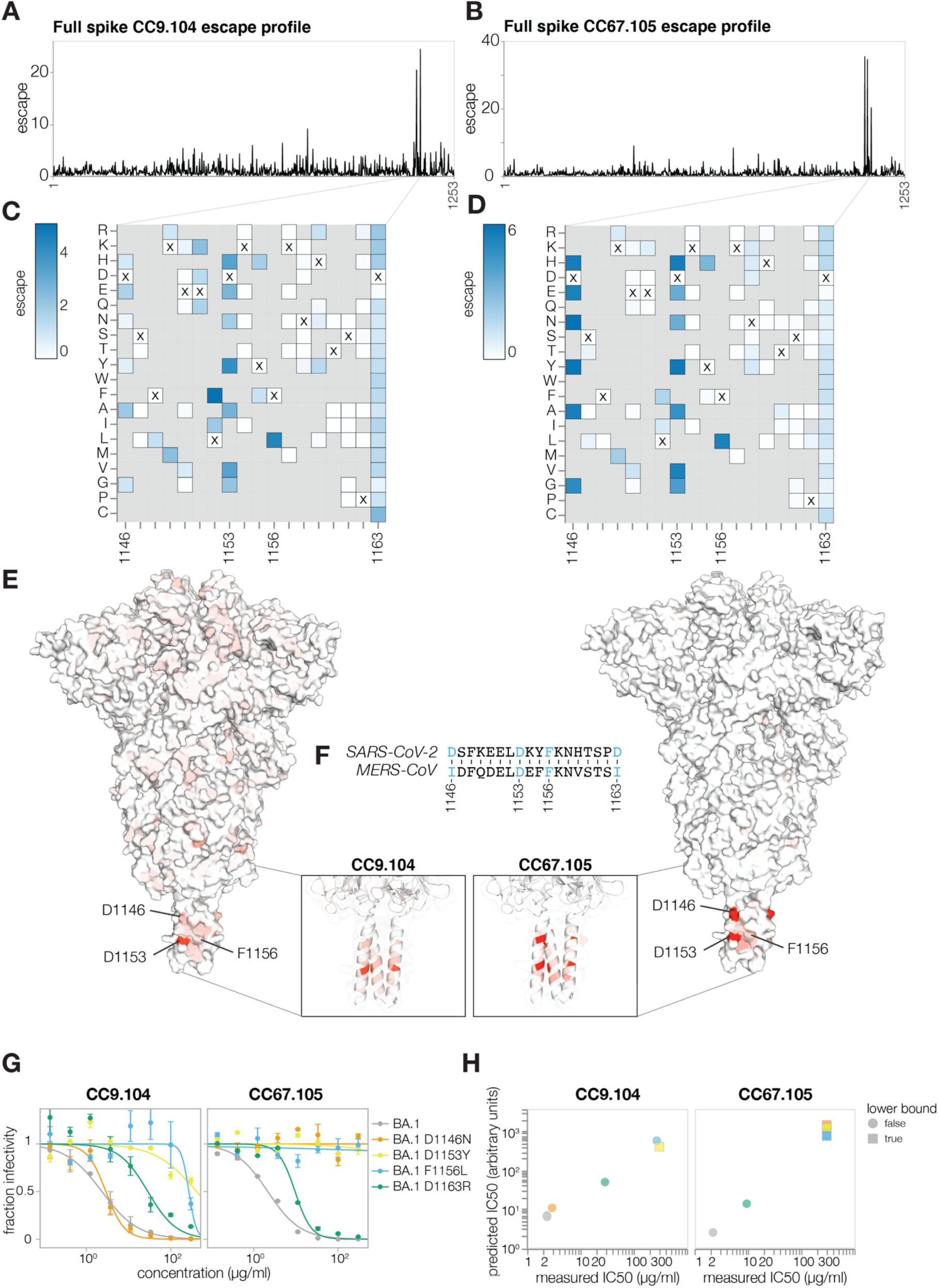
Antibody CC9.104 and CC67.105 escape mapping. **(A-B)** Total escape scores for each site in the BA. 1 spike for the CC9.104 (A) and CC67.105 (B) antibodies. **(C-D)** Escape heatmaps for the S2 stem-helix (sites 1146-1163) for CC9.104 (C) and CC67.105 (D) antibodies. Residues marked with X are the wild-type amino acids in BA.1 sequence. Amino acids that are not present in our libraries are shown in gray. Interactive heatmaps for the entirety of spike are at https://dms-vep.github.io/SARS-CoV-2_Omicron_BA.1_spike_DMS_mAbs/CC67.105_escape_plot.html and https://dms-vep.github.io/SARS-CoV-2_Omicron_BA.1_spike_DMS_mAbs/CC9.104_escape_plot.html **(E)** Surface representation of spike coloured by the sum of escape scores at that site for CC9.104 (left) and CC67.105 (right) antibodies. Site 1163 is not resolved in the structure. PDB ID: 6XR8. **(F)** Alignment of SARS-CoV-2 and MERS-CoV spikes at sites 1146-1163. **(G)** Validation pseudovirus neutralization assay for CC9.104 (left) and CC67.105 (right) antibodies with BA.1 spike carrying the indicated mutations. **(H)** Correlation between predicted IC50 values from deep mutational scanning (DMS) data versus the IC50 values measured in the validation assays in (G). Lower bound indicates that the antibody did not neutralize at the highest concentration tested in the validation neutralization assay. Site numbering in all plots is based on the Wuhan-Hu-1 sequence.

The above deep mutational scanning of escape from the S2 antibodies emphasizes the difference between SARS-related coronavirus breadth and resistance to escape in SARS-CoV-2. Both CC9.104 and CC67.105 neutralize many diverse SARS-related coronaviruses, but Omicron subvariants with mutations that lead to almost complete escape from these antibodies have been detected (e.g. D1153Y in BA.2.46 and BA.2.59). Therefore, even pan-sarbecovirus neutralizing antibodies can be escaped by mutational diversity within SARS-CoV-2, which emphasizes the importance of directly mapping escape mutations in SARS-CoV-2 in addition to assessing breadth across other natural SARS-related coronaviruses.

To show that we can perform deep mutational scanning of the spikes from different SARS-CoV-2 strains, we mapped escape from the REGN10933 antibody using Delta spike deep mutational scanning libraries (Figure S6A-B). REGN10933 is a class 1 antibody that directly competes with ACE2 binding, and was part of REGN-COV2 therapeutic cocktail used early in the pandemic but has lost potency against Omicron variants (Baum et al., 2020; Hansen et al., 2020; Liu et al., 2022). Escape sites for REGN10933 mapped with our deep mutational scanning system overlapped with the antibody binding footprint and included previously described escape mutations (Figure S6C) (Baum et al., 2020; Hansen et al., 2020; Starr et al., 2021b).

### Functional effects of mutations on spike-mediated pseudovirus infection

Our deep mutational scanning also enables measurement of how mutations affect spike-mediated viral infection in the absence of antibodies. We can make these measurements by computing a functional score for each variant from its relative frequency in infectious spike-pseudotyped lentiviruses generated from our single-copy cell integrated cells versus VSV-G-pseudotyped lentivirus generated from the same cells (Figures 1B). Spike variants with negative functional scores are worse at mediating cellular infection than the parental unmutated spike, while variants with positive functional scores are better at mediating infection (Figure 2). To deconvolve the functional scores for the variants (which often contain multiple mutations) into the effects of individual mutations on spike-mediated entry, we used global epistasis models (Otwinowski et al., 2018; Sailer and Harms, 2017).

As expected stop-codon mutations to the BA.1 spike were highly deleterious for spike-mediated infection, whereas amino-acid mutations showed a wide range of effects ranging from slightly beneficial to roughly neutral to highly deleterious (Figure 7A; recall that our library design excludes many of the most deleterious amino-acid mutations). To test whether the mutations measured to have slightly beneficial effects actually improved spike-mediated infection, we chose five mutations that the deep mutational scanning indicated improved infection (Figure 7B), and generated pseudovirus mutants carrying these mutations. The validation experiments confirmed that all the tested mutations indeed slightly improved spike-mediated infection (Figure 7C), validating that our deep mutational scanning can identify mutations that increase spike-mediated pseudovirus infection.

**Figure 7.**
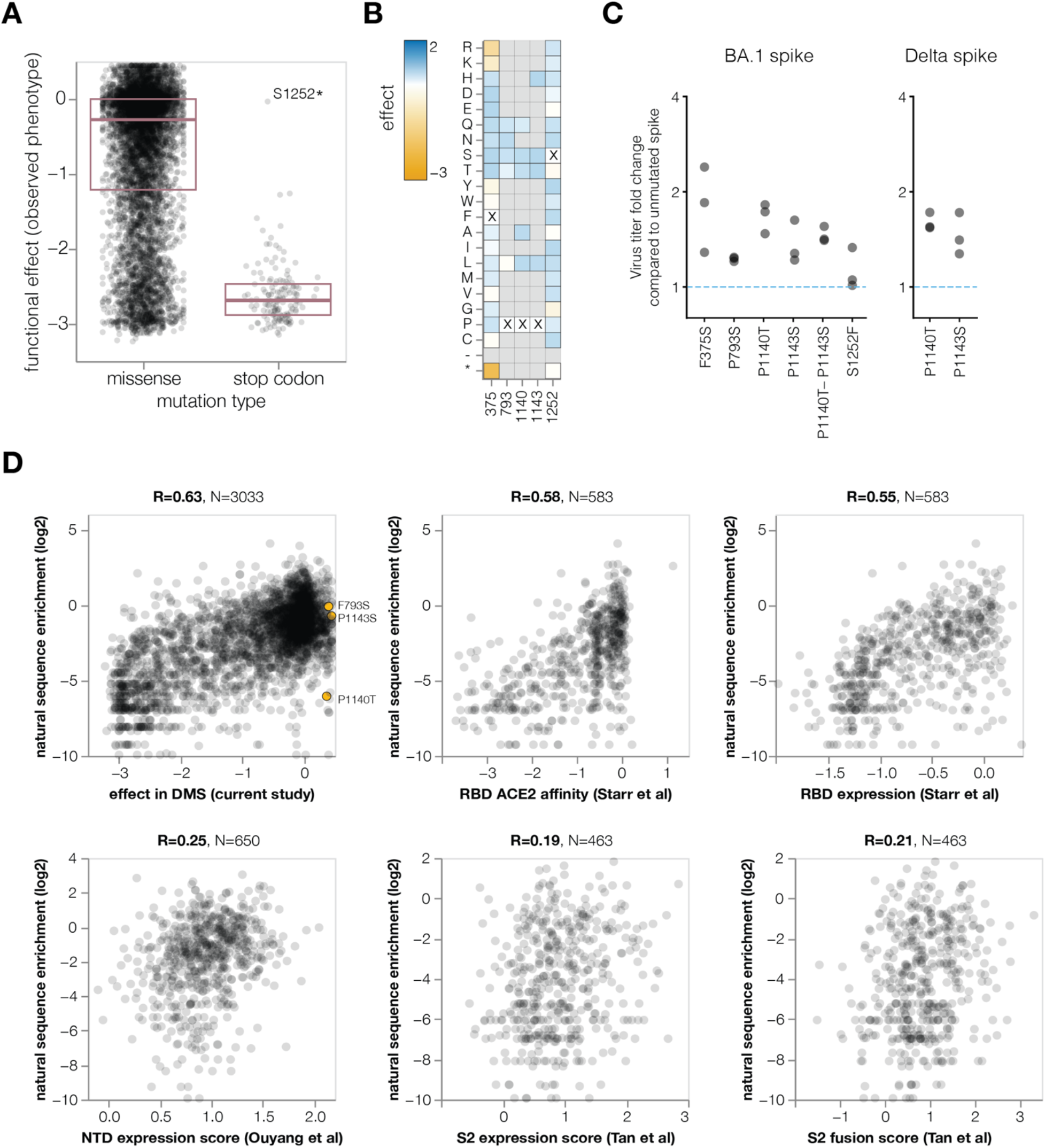
Functional effects of mutations on spike-mediated pseudovirus infection. **(A)** Distribution of functional effects of mutations in BA.1 deep mutational scanning libraries. Negative values indicate mutations are deleterious for viral entry. The stop codon mutation with a neutral functional effect of ~0 is at the last codon of the spike used in our experiments. **(B)** Heatmap showing functional effects at sites of mutations with beneficial functional effects that were chosen for validation assays in C. An interactive version of this heatmap for the entire spike is at https://dms-vep.github.io/SARS-CoV-2_Omicron_BA.1_spike_DMS_mAbs/muteffects_observed_heatmap.html **(C)** Fold change in virus entry titer for spike mutants relative to unmutated spike. There are three points for each mutant, reflecting triplicate measurements. **(D)** Correlation between enrichment of mutations during actual evolution of human SARS-CoV-2 and functional effects from our lentivirus-based deep mutational scanning or previous RBD expression or ACE2 affinity for yeast-based deep mutational scanning (Starr et al., 2022), and S2 (Tan et al., 2022) or NTD (Ouyang et al., 2022) expression for mammalian display-based deep mutational scanning. Interactive plots that enable mouseovers and show correlations among experiments are at https://dms-vep.github.io/SARS-CoV-2_Omicron_BA.1_spike_DMS_mAbs/natural_enrichment_vs_dms.html

To examine the relationship between the functional effects of spike mutations in the deep mutational scanning and actual evolution of human SARS-CoV-2, we determined the extent that mutations are enriched or depleted across a phylogenetic tree of all publicly available human SARS-CoV-2 sequences (Turakhia et al., 2021). To do this, we calculated the number of independent observations of each mutation on the tree and compared these observed numbers to the expected numbers under neutrality as estimated from four-fold synonymous sites, analyzing only mutations expected to have ≥20 occurrences (see Methods for details). Our deep mutational scanning measurements of the effects of mutations on spike-mediated infection were reasonably correlated with the enrichment of mutations among actual sequences (Figure 7D), indicating our experiments at least partially reflect the functional selection actually shaping spike evolution. We performed a similar analysis for prior spike deep mutational scanning using yeast display of the RBD (Starr et al., 2022), or mammalian cell display of the NTD (Ouyang et al., 2022) or a region of S2 (Tan et al., 2022) (Figure 7D). Our pseudovirus-based spike deep mutational scanning measurements were more correlated with the enrichment of mutations during actual evolution than any of these prior cell-surface display deep mutational scanning studies, presumably because our experiments mimic the true biological function of spike better than cell-surface display experiments.

However, none of the mutations with positive deep mutational scanning functional scores that we validated to improve spike-mediated infection in a pseudovirus context are enriched during actual SARS-CoV-2 evolution (Figure 7D). We suggest that this is because there is some divergence between the selection pressure in our pseudovirus-based experiments and true natural selection on spike. For instance, mutations at sites P1140 and P1143, which are located at the beginning of the S2 stem-helix, could potentially destabilize the prefusion trimer leading to more rapid cell entry in a pseudovirus context but negatively affect spike stability in the context of actual human transmission. Nonetheless, our functional measurements still provide the most accurate large-scale measurements to date on the effects of mutations to spike, and should be useful for assessing which antibody-escape mutations are well enough tolerated to pose a plausible risk of emerging naturally. We also note that our experiments indicate that there are no further mutations to the BA.1 spike that improve pseudovirus titers to the same extent as the D614G mutation that fixed early in SARS-CoV-2’s evolution in humans (Benton et al., 2021; Plante et al., 2021; Zhang et al., 2021).

## Discussion

We have developed a new deep mutational scanning system for assessing the antigenic and functional effects of mutations in the SARS-CoV-2 spike. This deep mutational scanning system is the first to measure how mutations to the entirety of spike affect cellular infection, and therefore enables the mapping of escape from antibodies targeting any part of the spike. Furthermore, our system directly quantifies how mutations affect antibody neutralization, and we show that these measurements correlate well with traditional pseudovirus neutralization assays. We expect that the ability to directly measure neutralization as opposed to binding will be especially useful when applied to polyclonal sera, since the magnitude of how mutations affect neutralization versus binding can differ in a polyclonal context (Bellusci et al., 2022; Cao et al., 2022).

Using the new deep mutational scanning system, we have mapped mutations that escape monoclonal antibodies targeting the RBD, NTD, and S2 domains. Although we only characterized a few antibodies here, this work is a first step towards generating similar data for much larger antibody panels. For the RBD, such data has proven valuable for interpreting the antigenic phenotypes of the new SARS-CoV-2 variants constantly emerging in the human population (Cao et al., 2022; Greaney et al., 2022). The ability to generate comparable data for other spike domains should enable better antigenic assessment of new variants, which can help inform vaccine design and strain selection (DeGrace et al., 2022). In addition, our work on the S2 antibodies emphasizes the distinction between breadth on natural viruses and potential for escape within SARS-CoV-2: an antibody that cross-neutralizes MERS-CoV and all tested SARS-related viruses is still escaped by mutations already present in some SARS-CoV-2 variants. Therefore, direct prospective mapping of escape mutations (Starr et al., 2021b, 2021a) should be useful for informing development of therapeutic antibodies targeting even regions of spike conserved across most natural viruses.

We also used the deep mutational scanning system to measure how mutations in spike affect its ability to mediate pseudovirus infection. These measurements complement existing deep mutational scanning datasets on how spike mutations affect the molecular phenotypes of ACE2 affinity (Starr et al., 2022), membrane fusion (Tan et al., 2022), and cell-surface expression (Ouyang et al., 2022; Starr et al., 2022; Tan et al., 2022). None of these experimental measurements fully reflect how mutations affect actual viral fitness, which is an emergent property of many molecular phenotypes (Ballal et al., 2020; Harms and Thornton, 2013). However, understanding how mutations affect molecular phenotypes is an important step towards interpreting evolutionary outcomes, and our pseudovirus-based system currently provides the deep mutational scanning measurements that best reflect natural selection on the spike of human SARS-CoV-2.

The deep mutational scanning system we describe here can also be straightforwardly extended to any virus with an entry protein amenable to lentiviral pseudotyping. This set of viruses includes other coronaviruses, influenza viruses, filoviruses, arenaviruses, and henipaviruses—all of which have receptor-binding and fusion proteins for which lentiviral pseudotyping provides a safe way to study cellular infection and antibody neutralization without requiring direct work with the actual pathogenic virus (Huang et al., 2020; Khetawat and Broder, 2010; Kobinger et al., 2001; Larson et al., 2008; Medina et al., 2003). Deep mutational scanning of the entry proteins of all these viruses could provide valuable information for antigenic surveillance and vaccine design, since these proteins are the dominant target of neutralizing antibodies. However, we note that data generated by such a system could in principle be used to inform introduction of gain-of-function mutations into actual potential pandemic viral pathogens. We therefore suggest that advances in the high-throughput characterization of mutations to viral proteins should be coupled with thoughtful limits on any downstream experiments with actual replicating viruses (Inglesby et al., 2022) to ensure that safely generated information is used to benefit public health without creating new risks.

## Methods

### Design of lentiviral backbone and spike gene nucleotide sequence optimization

The structure of lentiviral backbone is shown in Figure 1A. The plasmid map of the lentivirus backbone containing BA.1 spike is at https://github.com/dms-vep/SARS-CoV-2_Omicron_BA.1_spike_DMS_mAbs/blob/main/library_design/reference_sequences/3282_pH2rU3_ForInd_Omicron_sinobiological_BA.1_B11529_Spiked21_T7_CMV_ZsGT2APurR.gb and the map for the Delta spike-containing backbone is at https://github.com/dms-vep/SARS-CoV-2_Delta_spike_DMS_REGN10933/blob/main/library_design/reference_sequences/pH2rU3_ForInd_sinobiolo_gical_617.2_Spiked21_CMV_ZsGT2APurR.gb. Note our BA.1 spike includes the EPE insertion after position 214 (Wuhan-Hu-1 numbering). The vector is based on a pHAGE2 lentiviral backbone in which we repaired the 3’ LTR sequence (OhAinle et al., 2018), which allows us to re-rescue the pseudovirus from the cells in which lentiviral backbones have been integrated. The lentiviral backbone is non-replicative unless helper plasmids (Gag/Pol (NR-52517), Tat1b (NR-52518), and Rev1b (NR-52519)) are also transfected into the cells containing this backbone. Expression of the spike gene in the lentivirus backbone is driven both by inducible TRE3G promoter and by Tat1b. TRE3G promoter is activated by addition of doxycycline in the presence of the reverse tetracycline transactivator (rtTA), which is endogenously expressed in HEK-293T-rtTA cells. The spike gene has been codon optimized and lacks 21 amino acids in its cytoplasmic tail. The cytoplasmic tail deletion has been previously shown to significantly increase pseudovirus titers (Crawford et al., 2021; Havranek et al., 2020). For spike sequence codon optimization we tested a large panel of optimized sequences and found that virus titers can vary between codon optimizations by as much as 100-fold. While we did observe some titer differences when virus was generated from just transfection reactions using lentiviral backbones and helper plasmids, these differences were even more noticeable when virus was generated from cells with integrated spike-carrying lentiviral backbones. Of the tested codon optimizations we found that the sequence optimized spike from SinoBiological (VG40609-UT) gave by far the best virus titers; we therefore based all variant sequences on the original SinoBiological optimization. In addition to the inducible promoter and spike gene, the backbone also has a CMV promoter that drives expression of the ZsGreen gene linked by a T2A linker to the puromycin resistance gene. ZsGreen is used as a reporter gene to detect pseudovirus infection and the puromycin resistance gene is used as a selection marker for cells with successfully integrated lentiviral backbones.

### Design of spike mutations to include in BA. 1 and Delta full spike deep mutational scanning libraries

We aimed to create a library with mutations that would result in mostly functional spike proteins and include the mutations likely to be important for the antigenic evolution of spike. To this end, for BA.1 and Delta deep mutational scanning libraries we included the following types mutations: (1) mutations (nonsynonymous changes and deletions) observed in spike sequences deposited on GISAID database, (2) mutations that reoccur in spike phylogeny independently multiple times, (3) all possible amino acid changes at sites in spike that show positive selection. Specifically for the BA.1 library, we also included all possible amino acid changes for sites that are mutated in the BA.1 spike relative to Wuhan-Hu-1.

The following criteria were used to select the above described mutations for BA.1 library: nonsynonymous mutations need to be present in GISAID database >16 times, deletions need to occur in the NTD and be observed on GISAID database >300 times, nonsynonymous mutations need to reoccur on spike phylogenetic tree independently at least 21 times. To get all spike mutations observed in GISAID deposited sequences we used a CoVsurver curated spike amino acid frequency table (with sequences deposited up to January-31-2022) (Khare et al., 2021). To get independently recurring spike mutation counts we used pre-built SARS-CoV-2 phylogenies from UShER (Turakhia et al., 2021). Information on sites in spike undergoing positive selection was taken from taken from table here https://raw.githubusercontent.com/spond/SARS-CoV-2-variation/master/windowed-sites-fel-2021-07.csv which was built using methods described in Maher et al. (2022). The full list of mutations included in the BA.1 library can be found at https://github.com/dms-vep/SARS-CoV-2_Omicron_BA.1_spike_DMS_mAbs/blob/main/library_design/results/aggregated_mutations.csv.

The following criteria were used to select the above described mutations for the Delta library: nonsynonymous mutations and deletions need to be observed on GISAID database more than once and nonsynonymous mutations need to reoccur on spike phylogenetic tree independently more than 7 times. To get all spike mutations observed in GISAID deposited sequences we aligned all spike sequences deposited on GISAID up to July-26-2021 and extracted mutation frequency counts. Independently recurring spike mutations and positively selected sites were identified as described for BA.1 library above. The full list of mutations included in the Delta library can be found at https://github.com/dms-vep/SARS-CoV-2_Delta_spike_DMS_REGN10933/blob/main/library_design/results/aggregated_mutations.csv.

### Design of primers for BA.1 and Delta spike mutagenesis

For each set of mutations described in the section above we designed separate primer pools: (1) a pool of primers for observed mutations, (2) a pool of primers for recurrent mutations, (3) a pool of primers for positive selection site mutations, and (4) a pool of primers that would cover changes at multiple positive selection sites if those positive selection sites are close enough to each other so that the primers in the pool (3) would overlap. For the BA.1 library we also designed primers that would introduce multiple amino acid deletions at recurrent deletion regions described in (McCarthy et al., 2021) and included them in the observed mutation primer pool. Also for the BA.1 library, the set of primers that cover all possible amino acid changes at the sites already mutated in BA.1 was pooled with the positive selection site primer pool.

All primer pools were ordered from Integrated DNA Technologies as oPools. Scripts for designing the BA.1 library primer pools and the resulting oPools that were ordered are at https://github.com/dms-vep/SARS-CoV-2_Omicron_BA.1_spike_DMS_mAbs/tree/main/library_design and scripts for designing the Delta library primer pools are at https://github.com/dms-vep/SARS-CoV-2_Delta_spike_DMS_REGN10933/tree/main/library_design.

### Design of full-spike deep mutational scanning plasmid libraries

Making of the plasmid libraries for deep mutational scanning required the following three steps (1) mutagenesis of the spike gene, (2) barcoding of the mutagenised spike sequence, and (3) cloning of the mutagenised and barcoded spike into the lentiviral backbone-carrying plasmid.

Spike mutagenesis was carried out by first amplifying BA.1 or Delta spike gene sequence from a plasmid carrying lentiviral backbone with a codon optimized spike sequence (see section ‘Design of lentiviral backbone and spike gene nucleotide sequence optimization’ for plasmid maps). Spike sequence was amplified using ‘Spike amplification’ primers from Supplementary Table 1 with the following PCR conditions: 1.5 μl of 10μM forward primer, 1.5 μl of 10 μM reverse primer, 10 ng of amplified spike gene template, 25 μl of KOD polymerase (KOD Hot Start Master Mix, Sigma-Aldrich, Cat. No. 71842), and water for the final volume of 50 μl. PCR cycling conditions were as follows:

1. 95°C for 2 min
2. 95°C for 20 s
3. 62°C for 15 s
4. 70°C for 2 min (return to step 2 for another 19x cycles)
5. Hold at 4°C

Amplified spike sequence was first gel-purified using NucleoSpin Gel and PCR Clean-up kit (Takara, Cat. No. 740609.5) and then further purified using Ampure XP beads (Beckman Coulter, Cat. No. A63881) at 1:2.6 sample to bead ratio.

Next the purified spike template was used in mutagenesis PCR using protocol described previously in (Bloom, 2014) with a few modifications (see also https://github.com/jbloomlab/CodonTilingPrimers for general background on this protocol). Forward and reverse primers for mutagenesis PCR were pooled into separate pools at 1: 2: 2: 0.2 per primer molar ratio between observed primer pool: recurrent primer pool: positive selection primer pool: paired positive selection primer pool. The pooling ratios are determined by the fact that recurrent and positively selected sites may be more antigenically and structurally important for spike. Two independent mutagenesis reactions were performed for each spike creating two independent biological library replicates (which means that they will have a unique set of barcodes and a unique set of mutation combinations in spike). For BA.1 libraries we performed two rounds of mutagenesis with the first round consisting of 8 mutagenic PCR cycles followed by the second round of 10 mutagenic PCR cycles. For Delta libraries one biological replicate consisted of a single round of 10 mutagenic PCR cycles and the second biological replicate consisted one round of 8 and another round of 10 mutagenic PCR cycles. Each mutagenesis PCR reaction was divided into forward and reverse reactions based on whether the reaction used the forward or reverse pool of mutagenesis primers. The PCR conditions for mutagenesis were as follows: 1.5 μl of 5 μM of forward or reverse mutagenesis primer pool, 1.5 μl of 5 μM of forward or reverse ‘Spike amplification and joining PCR’ primer (see supplementary table 1), 1.5 μl of 3 ng/μl linearised spike template, 4 μl of water, and 15 μl of KOD. The PCR cycling conditions were as follows:

1. 95°C, 2 min
2. 95°C, 20 s
3. 70°C, 1 s
4. 54°C, 20 s, cooling at 0.5°C/s
5. 70°C, 100 s (return to step 2 for the number of cycles described above)
6. 4°C, hold

Between each mutagenic PCR round 20 cycles of joining PCR were performed. For joining PCR we used 4 μl each of 1:4 diluted forward and reverse mutagenesis reactions, 1.5 μL each of forward and reverse joining primers (see ‘Spike amplification and joining PCR’ primers in supplementary table 1), 4 μl of water, and 15 μl of KOD. PCR cycling conditions were identical to the mutagenesis PCR conditions. Joined PCR mutagenesis products were gel and Ampure XP purified after each joining reaction.

After the spike sequence was mutagenised we performed a barcoding PCR that appended a random 16 nucleotide barcode sequence downstream of the spike gene stop codon. We chose 16 nucleotide barcodes as it allows for a total of 4^16^ unique barcoded variants, which is much greater diversity of barcodes than the final size of our deep mutational scanning plasmid libraries and therefore limits potential barcodes dulications. For barcoding ‘Spike barcoding’ primers from supplementary table 1 were used with the following PCR conditions 1.5 μl of 10μM forward primer, 1.5 μl of 10 μM reverse primer, 30 ng of the mutagenised spike gene template, 25 μl of KOD polymerase, and water for final volume of 50 μl. PCR cycling conditions were as follows:

1. 95°C, 2 min
2. 95°C, 20 s
3. 70°C, 1 s
4. 55.5°C, 20 s, cooling at 0.5°C/s
5. 70°C, 2 min (return to step 2 for another 9x cycles)
6. 4°C hold

The mutagenised and barcoded spike was then cloned into lentiviral backbone-containing plasmid. First, we digested a lentiviral backbone containing plasmid using Mlul and Xbal restriction sites. The map of plasmid used for vector digestion is at https://github.com/dms-vep/SARS-CoV-2_Omicron_BA.1_spike_DMS_mAbs/blob/main/library_design/reference_sequences/other_plasmid_maps_f_or_library_design/3137_pH2rU3_Forlnd_mCherry_CMV_ZsGT2APurR.gb. Digested vector was gel and Ampure XP purified. We then used 1:3 insert to vector ratio in a 1 hour Hifi assembly reaction using NEBuilder HiFi DNA Assembly kit (NEB, Cat. No. E2621). After the HiFi assembly, we Ampure XP purified the reaction and eluted it in 20 μl of water (note that elution in water as opposed to elution buffer enhances the subsequent electroporation efficiency). We used 1 μl of the purified HiFi product to transform 20 μl of 10-beta electrocompetent *E. coli* cells (NEB, C3020K). We performed 10 electroporation reactions to get a final count of > 2 million CFUs per library and plated transformed cells out on LB+ampicillin plates. We aim to make plasmid library from a much greater number of CFUs than the number of variants in our final virus libraries to minimize barcode duplication, as explained in the next section. About 16 hours after transformation bacterial colonies were scraped using liquid LB+ampicillin and plasmid stocks were prepared using QIAGEN HiSpeed Plasmid Maxi Kit (Cat. No. 12662). The final structure of the lentiviral genome with mutagenised spike cloned into it is shown in Figure 1A.

### Production of cell-stored spike deep mutational scanning libraries

Production of cell-stored deep mutational scanning libraries required the following steps: (1) production of VSV-G-pseudotyped lentivirus, (2) infection of rtTA-expressing cells with VSV-G-pseudotyped virus, and (3) selection for transduced cells. These steps are illustrated in Figure 1B.

To generate VSV-G-pseudotyped virus for each library we plated 0.5 million HEK-293T cells per well in eight wells of two 6-well tissue culture dishes. Note we aim to produce VSV-G-pseudotyped virus stocks that have a greater number of infectious particles than the number of colonies scraped for plasmid libraries in order to not introduce any bottleneck on barcodes at this stage. The next day we transfected 0.25 μg of each helper plasmid (Gag/Pol, Tat1b, and Rev1b), 0.25 μg of VSV-G expression plasmid (https://github.com/jbloomlab/SARS-CoV-2-BA.1_Spike_DMS_validations/blob/main/plasmid_maps/29_HDM_VSV_G.gb) and 1 μg of mutagenised and barcoded spike containing lentiviral vector (described in the section above). Transfections were done using BioT reagent (Bioland Scientific, Cat. No. B01-02) according to the manufacturer’s instructions. 48 hours post transfection supernatants from each well were pooled, filtered through a surfactant-free cellulose acetate 0.45 μm syringe filter (Corning, Cat. No. 431220), and stored at −80°C. VSV-G-pseudotyped viruses were titrated as described in (Crawford et al., 2020).

Next we infected HEK-293T-rtTA cells with the generated VSV-G-pseudotyped virus. The number of infectious virus units used in these infections allowed us to bottleneck the library size at the desired final variant number. For BA.1 libraries we attempted to bottleneck libraries at 100,000 variants and for Delta libraries we bottlenecked the libraries at 50,000 variants. Notably, we used a substantially lower number of variants to infect cells compared to the possible diversity of variants in our plasmid libraries. This allows us to limit any potential duplication of barcodes between different variants due to recombination in the lentivirus genome (Hill et al., 2018; Jetzt et al., 2000; Schlub et al., 2010). Note for BA.1, libraries Lib-1 and Lib-2 originate from the same mutagenised lentiviral backbone plasmid stock but independent VSV-G virus infections and Lib-3 originates from independent mutagenised plasmid library stock. For Delta libraries Lib-1 and Lib-2 are both from independent mutagenised spike plasmid stocks. Infections were performed at MOI < 0.01 (in order to ensure that only a single spike variant is integrated in each cell), which was verified 48 hours after infection using fluorescence-activated cell sorting by detecting ZsGreen expression from the lentiviral backbone. After MOI was verified, we expanded cells for another 48 hours and then started puromycin selections to select for cells with successfully integrated lentivirus genomes. Selection was done using 0.75 μg/ml of puromycin with a fresh change of puromycin-containing D10 (see ‘Cell lines’ section below) every 48 hours. Selections were terminated when visual inspection using a fluorescent microscope indicated that all cells express ZsGreen (approximately 6-8 days). After puromycin selection was finished we expanded cells for another 48 hours in fresh D10 and froze cell aliquots in tetracycline-free FBS (Gemini Bio, Cat. No. 100-800) containing 10% DMSO. Frozen cell aliquots were stored in liquid nitrogen long-term.

### Generation of spike and VSV-G-pseudotyped viruses from cell-stored spike deep mutational scanning libraries

To generate spike-pseudotyped viruses from cell-stored deep mutational scanning libraries we plated 100 million library-containing cells per 5-layer flask (Corning Falcon 875cm^2^ Rectangular Straight Neck Cell Culture Multi-Flask, Cat. No. 353144) in 150 ml of D10 without phenol red supplemented with 1 μg/ml for doxycycline (which allows to induce spike expression ahead of pseudovirus production). 24 hours after plating we transfected cells with 50 μg of each helper plasmid (Gag/Pol, Tat1b, Rev1b) using BioT reagent according to manufacturer’s instructions. 48 hours post transfection cell supernatant was collected and filtered through a 0.45 μm SFCA Nalgene 500mL Rapid-Flow filter unit (Cat. No. 09-740-44B). Filtered supernatant was then concentrated by spinning at 4°C 3000 rcf for 30 min using Pierce Protein Concentrator (ThermoFisher, 88537). Virus aliquots were stored long-term at −80°C. Titers for concentrated spike-pseudotyped libraries titrated on HEK-293T-ACE2 cells ranged between 0.5×10^6^ - 2×10^6^ transcription units per ml.

To generate VSV-G-pseudotyped viruses (for functional selection and long-read PacBio sequencing) from cell-stored deep mutational scanning libraries we plated 60 million library-containing cells per 3-layer flask (Corning Falcon 525cm2 Rectangular Straight Neck Cell Culture Multi-Flask, 353143) in 90 ml of D10 without phenol red (note we do not add doxycycline in this case). 24 hours after plating we transfected cells with 30 μg of each of the helper plasmid (Gag/Pol, Tat1b, Rev1b) and 18.75 μg of VSV-G expression plasmid using BioT reagent according to manufacturer’s instructions. 32-36 hours post transfection cell culture supernatant was collected and filtered through a 0.45 μm SFCA Nalgene filter unit. Filtered supernatant was then concentrated by spinning at 4°C 3000 rcf for 30 min using Pierce Protein Concentrator. Virus aliquots were stored long-term at −80°C. Titers for concentrated VSV-G-pseudotyped libraries titrated on HEK-293T-ACE2 cells ranged between 10×10^6^ - 30×10^6^ transcription units per ml.

### Long-read PacBio sequencing of barcoded spike variants in deep mutational scanning libraries

Long-read PacBio sequencing was used to acquire reads spanning the spike and the random 16 nucleotide barcode sequences. To prepare amplicons for PacBio sequencing we infected 1 million HEK-293T cells with ~30 million VSV-G-pseudotyped lentiviruses carrying the deep mutational scanning libraries. This number of viruses is significantly greater than the expected number of variants in the library, which allows us to achieve high variant coverage, avoid bottleneck of barcode diversity and correct for any potential PCR or sequencing errors. 12-15 hours after infection cells were trypsinized, washed with PBS and non-integrated lentiviral genomes were recovered using QIAprep Spin Miniprep Kit (Cat. No. 27106X4) (Dingens et al., 2018; Haddox et al., 2016). We use non-integrated viral genomes as our sequencing templates because they are the more abundant forms of the lentiviral genome than the integrated proviruses (Chun et al., 1997; Pang et al., 1990; Sharkey et al., 2000; Van Maele et al., 2003). Elution volume for the miniprep was adjusted to 144 μl. Next we performed two rounds of PCR to amplify the region in the lentivirus genome spanning the spike and the random 16 nucleotide barcode. In the first round of PCR we use primers containing single nucleotide tags, which allow us to later detect strand exchange that may occur during PCR amplification. To limit strand exchange during PCR (which would disrupt barcode/spike variant linkage) we also minimize the number of PCR cycles performed and do multiple PCR reactions per sample (Liu et al., 2014; Omelina et al., 2019) . Each sample was split into eight PCR reactions, four of which use ‘tag_1’ forward and reverse primers and four of which use ‘tag_2’ forward and reverse primers from the ‘Spike gene amplification for PacBio long-read sequencing’ primer set in Supplemental Table 1. PCR reaction conditions were as follows: 1 μl of forward primer, 1 μl of reverse primer, 20 μl of KOD, and 18 μl of sample. PCR cycling conditions for round 1 PCR were as follows:

1. 95°C for 2 min
2. 95°C for 20 s
3. 70°C for 1 s
4. 60°C for 10 s (ramp 0.5°C/s)
5. 70°C for 2.5 min (go to 2 for another 7 cycles)
6. 70°C for 5 min
7. 4°C hold

After the first PCR round we pooled all reactions for each sample and purified them using Ampure XP beads with 1:0.8 beads to sample ratio and the PCR product was eluted in 84 μl of elution buffer. Eluted PCR product was divided into four PCR tubes and the second round of PCR was performed using ‘RND2’ forward and reverse primers from the ‘Spike gene amplification for PacBio long-read sequencing’ primer set in the Supplemental Table 1. PCR reaction conditions were as follows: 2 μl of forward primer, 2 μl of reverse primer, 25 μl of KOD, and 21 μl of purified sample. PCR cycling conditions were the same as for the round 1 PCR for a total of 10 PCR cycles. PCR reactions for each sample were pooled, purified using Ampure XP beads with 1:0.8 beads to sample ratio, and eluted in 27 μl of elution buffer. Barcodes were attached to each sample using sample SMRTbell prep kit 3.0 before multiplexing. Multiplexed SMRTbell libraries were then bound to polymerase using Sequel II Binding Kit 3.2 and sequenced with PacBio Sequel IIe sequencer with a 20-hour movie collection time.

### Antibody escape mapping using full spike deep mutational scanning libraries

For antibody escape mapping we used between 4-15 times more infectious virions than the estimated total number of barcodes in a deep mutational scanning library. Using significantly more infectious virions relative to the number of variants per library avoids bottlenecking by having multiple copies of each variant. Note that we expect there to be several fold more lentiviral genomes per selection experiment than the amount of infectious units used because we are recovering the non-integrated viral genomes for sequencing, which are more abundant than integrated proviral DNA (Chun et al., 1997; Pang et al., 1990; Sharkey et al., 2000; Van Maele et al., 2003) on which our library virus titers are based. For each antibody escape mapping experiment we made a master mix of library spike-pseudotyped virus mixed with VSV-G pseudotyped neutralization standard (described below). Neutralization standard was added at 1-2% of the total virus titer used in the experiment. Virus master mix was then aliquoted into eppendorf tubes to which either different mounts of antibody or no antibody was added. For LY-CoV1404, CC9.104 and CC67.105 antibodies we performed selection experiments at 3 concentrations, starting with IC_99_ concentration predetermined using standard pseudovirus neutralization assay and then increasing this concentration 4 fold and 16 fold. We start with IC_99_ concentration intending that around 1% of the library will be able to escape antibody selection. We use additional concentrations as it helps us to cover a greater dynamic concentration range in cases where the exact IC_99_ value is difficult to determine. Also, the use of multiple concentrations enables more precise mutation-escape predictions by the biophysical model used to decompose single-mutation effects (Yu et al., 2022). For LY-CoV1404 starting concentration was 0.654 μg/ml, for CC9.104 - 68 μg/ml, for CC67.105 - 52.5 μg/ml. For the REGN10933 we started at IC_99.5_ at 0.146 μg/ml and also increased that concentration by 4 fold and 16 fold. For the NTD 5-7 antibody, which does not fully neutralize the virus, we started with >IC96 concentration at 150 μg/ml and then increased that concentration by 2 fold. Virus was mixed with the antibody by inverting tubes several times, spun down at 300 g, and incubated at 37°C for 1 h. After incubation virus and antibody mix or no antibody control were used to infect approximately 0.5 million target cells, which were plated a day before in D10 supplemented with 2.5 μg/ml of amphotericin B (Sigma, Cat. No. A2942) (which increases viral titers as shown in Figure S1B). The target cell line for different antibodies is determined by whether an antibody is able to neutralize pseudovirus on that cell line. We have previously described this phenomena in (Farrell et al., 2022) where we show that non-ACE2 competing antibodies do not fully neutralize pseudovirus on ACE2 overexpressing cells. While testing antibodies for the current study we also noticed that some S2-targeting antibodies are also not affected by ACE2 overexpression. Therefore, for LY-CoV1404, CC9.104 and CC67.105 antibodies we used HEK-293T-ACE2 as target cells but for NTD-targeting 5-7 antibody we used HEK-293T-ACE2-medium cells. For REGN10933 we used HEK-293T-ACE2-TMPRSS2 as target cells, because TMPRSS2 overexpression increases Delta pseudovirus titers. 12-15 hours after infection cells were trypsinized, washed with PBS and non-integrated lentiviral genomes were recovered using QIAprep Spin Miniprep Kit and eluted in 21 μl of Qiagen elution buffer. Barcode reads for each sample were then prepared for Illumina sequencing using a method described in ‘Barcode amplicon preparation for Illumina sequencing’ section below.

### Functional selections using full spike deep mutational scanning libraries

To perform functional spike selections we infected 1 million HEK-293T-ACE2 cells with 1-2 millions of the spike or VSV-G-pseudotyped viruses produced from deep mutational scanning library carrying cells (described earlier). As for antibody selections, the amount of virus used is greater than the number of variants in each library which limits potential bottlenecking of the library barcodes. 12-15 hours after infection cells were trypsinized, washed with PBS and non-integrated lentiviral genomes were recovered using QIAprep Spin Miniprep Kit. Barcode reads for each sample were then prepared for Illumina sequencing using methods described in ‘Barcode amplicon preparation for Illumina sequencing’ section below.

### Barcode amplicon preparation for Illumina sequencing

To prepare barcode reads for Illumina sequencing we performed two rounds of PCR. In the first round of PCR we used primers that align to Illumina Truseq Read 1 primer site located directly upstream of the barcode in the lentiviral backbone and a primer annealing downstream of the barcode containing an overhand with Illumina Truseq Read 2 sequence (see ‘Illumina barcode sequencing 1st round PCR primers’ in the Supplemental Table 1). Conditions for the first round PCR were as follows: 1 μl of 10uM forward primer, 1 μl of 10uM reverse primer, 26 μl of KOD, and 24 μl of miniprepped sample DNA. PCR cycling conditions for round 1 PCR were as follows:

1. 95°C for 2 min
2. 95°C for 20 s
3. 70°C fro 1 s
4. 58°C for 10 s, cooling at 0.5°C per s
5. 70°C 20 s (return to step 2 for another 27 cycles)
6. 4°C hold

PCR reactions were purified with Ampure XP beads using 1:3 sample to beads ratio and eluted in 37 μl of Qiagen elution buffer. Second round of PCR used primers primer annealing to the Illumina Truseq Read 1 primer site with P5 Illumina adapter overhang and reverse primers from the PerkinElmer NextFlex DNA Barcode adaptor set, which anneal to Truseq Read 2 site and contain P7 Illumina adapter and i7 sample index. Conditions for the second round PCR were as follows: 1.5 μl of 10uM universal primer, 1.5 μl of 10uM indexing primer, 25 μl of KOD, and 20 ng of first round PCR product. PCR cycling conditions were the same as the first round PCR for a total of 20 cycles. After the second PCR round all samples were pooled at desired ratios and gel and Ampure XP bead purified. Barcode amplicons were sequenced using NextSeq 2000 with either P2 or P3 reagent kits.

### Production of barcoded neutralization standard

To make the neutralization standard that we add into our deep mutational scanning libraries we used the same general barcodinging approach as described above for the deep mutational scanning plasmid library generation with a few important differences. The lentiviral backbone used for the neutralization standard consists of TRE3G inducible mCherry protein and CMV promoter driven ZsGreen. The plasmid map of the template backbone is at https://github.com/dms-vep/SARS-CoV-2_Omicron_BA.1_spike_DMS_mAbs/blob/main/library_design/reference_sequences/other_plasmid_maps_for_library_design/2871_pH2rU3_ForInd_mCherry_CMV_ZsG_NoBC_cloningvector.gb. Note this backbone does not encode any viral glycoproteins and to rescue VSV-G-pseudotyped virus we provide VSV-G expression plasmid in *trans*. We amplified mCherry plasmid from the lentiviral template and barcoded it in two independent PCR reactions using 2 sets of primers containing 4 unique barcodes (see ‘Neutralization standard barcoding primers’ in supplementary table 1). Importantly, the unique barcodes were balanced so that there is a unique nucleotide at each position of the 16-nucleotide barcode between each of the four barcoding primers in a PCR reaction. Furthermore, the 8 barcoding primers are unique to the neutralization standard and are not present in any of our deep mutational scanning libraries. The PCR for barcoding was done the same way as described for deep mutational scanning plasmid library production and both PCR reactions were pooled together before HIfi assembly into the lentiviral backbone. VSV-G-pseudotyped lentiviral particles were then produced as described above (Production of cell-stored spike deep mutational scanning libraries) for the spike libraries, except using this mCherry-containing barcoded backbone as the lentiviral backbone. This VSV-G-pseudotyped virus was then used to infect HEK-293T-rtTA cells at low MOI. Successfully transduced HEK-293T-rtTA were then selected by flow activated fluorescence sorting based on ZsGreen expression and expanded. We then generated VSV-G-pseudotyped neutralization standard by tranfecting helper plasmids and VSV-G expression plasmid in the same way as described for deep mutational scanning library virus rescues. Note, that we use the neutralization standard generated from the integrated cells as opposed to the original transfection in order to prevent lentiviral backbone-containing plasmid contamination of the virus stocks that can occur when viruses are produced from transfections.

### Validation of deep mutational scanning by pseudovirus titration and neutralization

Spike genes carrying desired mutations were cloned by performing PCR reactions with partially overlapping desired mutation-containing primers followed by HiFi assembly. HDM_omicron_B11529_IDTDNA plasmid was used as the template for PCR. The map of the plasmid can be found at https://github.com/jbloomlab/SARS-CoV-2-BA.1_Spike_DMS_validations/blob/main/plasmid_maps/3277_HDM_omicron_B11529_IDTDNA.gb. All plasmid sequences were verified using full plasmid sequencing by Primordium. Mutated spike plasmids or VSV-G expression plasmid were then used to generate and titrate pseudoviruses as described in (Crawford et al., 2020) except that the backbone used for virus generation was pHAGE6_Luciferase_IRES_ZsGreen and which also only required Gag/Pol helper plasmid for virus rescues. Note, for the spike variants cloned to validate functional selections we performed three replicate virus rescues for each variant and each rescue was done using an independent plasmid preparation for that spike variant.

BA.1 spike variants rescued for funcional selection validation were titrated on HEK-293T-ACE2 and Delta spike variants were titrated on HEK-293T-ACE2-TMPRSS2 cells. We performed duplicate serial dilutions using supernatants collected from the virus rescues and measured luciferase expression at each dilution using Bright-Glo Luciferase Assay System (Promega, E2610). Virus titers were calculated as relative light units (RLU) per μl for each dilution and taking the average RLU/μl values across dilutions within a linear range. For spike variants used to validate antibody escape experiments virus titration was performed in the same way using the same target cells as the neutralization assays were performed in (see below).

For pseudovirus neutralization 12.5 thousand target cells were plated into poly-L-lysine coated, black-walled, 96-well plates (Greiner 655930) in D10 supplemented with 2.5 μg/ml of amphotericin B. For neutralization assays using LY-CoV1404, CC9.104 or CC67.105 antibodies we used HEK-293T-ACE2 as target cells, for REGN10933 we used HEK-293T-ACE2-TMPRSS2 as target cells, and for NTD 5-7 mAb we used HEK-293T-ACE2-medium as target cells. The use of different cell lines for each antibody is determined by the ability of that antibody to neutralize the virus on that cell line as described previously. Next day we prepared replicate serial dilutions for each antibody. Starting concentration for each antibody was as follows: LY-CoV1404 – 4 μg/ml, CC9.104 and CC67.105 – 300 μg/ml, 5-7 – 96 μg/ml, REGN10933 – 6 μg/ml. Serial dilutions were then mixed with pseudovirus and incubated for 1 h at 37°C. After incubation the virus-antibody mix was transferred onto the target cells. 48-55 h after infection Bright-Glo Luciferase Assay System (Promega, E2610) was used to measure luciferase activity. Fraction infectivity for each antibody dilution was calculated by subtracting background readings and dividing RLU values in the presence of antibody by RLU values in the absence of it. Neutralization curves were then plotted by fitting a Hill curve to fraction infectivity values using neutcurve software (https://jbloomlab.github.io/neutcurve/, version 0.5.7). Neutcurve package was also used to extract target IC_x_ values from the fitted neutralization curves.

Code for plotting virus titers and neutralization curves from this paper can be found at https://github.com/jbloomlab/SARS-CoV-2-BA.1 Spike DMS validations.

### Cell lines

HEK-293T were acquired from ATCC (CRL3216), HEK-293T-ACE2 cells are described in (Crawford et al., 2020), generation and characterization of HEK-293T-ACE2-medium cells is described in (Farrell et al., 2022) (referred to ‘medium’ cells in the reference), generation of HEK-293T-rtTA cells is described below. All cells were grown in D10 media (Dulbecco’s Modified Eagle Medium with 10% heat-inactivated fetal bovine serum, 2 mM l-glutamine, 100 U/mL penicillin, and 100 μg/mL streptomycin). For antibody selection experiments D10 was made with phenol-free DMEM (Corning DMEM With 4.5g/L Glucose, Sodium Pyruvate; Without L-Glutamine, Phenol Red from Fisher, Cat. No. MT17205CV). For experiments with HEK-293T-rtTA cells D10 was made with tetracycline-free FBS (Gemini Bio, Cat. No. 100-800) to avoid any expression of spike unless doxycycline is added.

To produce HEK-293T-rtTA expressing cells (used for storing deep mutational scanning libraries and required for TRE3G promoter activation) we first generated VSV-G-pseudotyped lentivirus carrying rtTA gene. To produce this virus we transfected 0.5 million HEK-293T cells with 0.25 μg of each helper plasmid (Gag/Pol, Tat1b, Rev1b), 0.25 μg of VSV-G expression plasmid and 1 μg of lentiviral backbone carrying plasmid into which rtTA has been cloned (plasmid map https://github.com/dms-vep/SARS-CoV-2_Omicron_BA.1_spike_DMS_mAbs/blob/main/library_design/reference_sequences/other_plasmid_maps_for_library_design/2850_pHAGE2_EF1a_TetOn_IRES_mCherry.gb). Note this lentiviral backbone is self-inactivating and therefore would not be produced and packaged into lentiviral particles upon helper plasmid transfection. 48 hours after transfection virus-containing cell supernatant was collected and filtered through a surfactant-free cellulose acetate 0.45 μm syringe filter. 5μl of the virus was used to infect 0.5 million low passage HEK-293T cells. 48 hours post infection single cell clones were sorted into a 96-well plate using BD Aria II cell sorter with 610/20 filter in PE-Texas Red channel. Single clones were expanded and tested for ability to produce high virus titers. High virus titers are essential for performing deep mutational scanning experiments on a practical experimental scale and we found that individual cell clones can vary significantly in the virus titers they can produce. Clonal cell population producing the highest virus titers was selected for expansion, frozen down in 10% DMSO and 20% FBS D10 media, and stored long-term in liquid nitrogen freezer.

### Antibodies

LY-CoV1404 antibody was cloned and produced by GensScript. Variable domain sequences were taken from previously published antibody structure (Westendorf et al., 2022). LY-CoV1404 variable regions were cloned with IgG1 heavy chain and human IgL2 constant regions, expressed in mammalian cells and purified using IgG-binding columns. S2 antibodies were produced as described previously in (Zhou et al., 2022). 5-7 antibody was produced as described previously in (Liu et al., 2020)

## COMPUTATIONAL ANALYSIS AND DATA AVAILABILITY

### Overview of data analysis pipeline

To analyze the deep mutational scanning data, we created a modular analysis pipeline. At the core of this pipeline is a set of common steps that are expected to be shared across analysis of many different datasets. We implemented this set of common steps in a standalone GitHub repository named *dms-vep-pipeline* that is publicly available at https://github.com/dms-vep/dms-vep-pipeline and is designed to be included in project-specific analyses as a git submodule. The *dms-vep-pipeline* consists of a series of Snakemake (Mölder et al., 2021) rules that run Python scripts or Jupyter notebooks, and specifies a conda environment that provides details on the software used for the analysis. For this paper, we used version 1.01 of the *dms-vep-pipeline*.

For each specific project (in this case, deep mutational scanning of the BA.1 and Delta spikes) we created a separate GitHub repository that included *dms-vep-pipeline* as a submodule. The repository for BA.1 is publicly available at https://github.com/dms-vep/SARS-CoV-2_Omicron BA.1 spike DMS mAbsand the repository for Delta is at https://github.com/dms-vep/SARS-CoV-2_Delta_spike_DMS_REGN10933 Note how each repository has a configuration file (the *config.yaml* file), project-specific input data (the *data* subdirectory), and a top-level Snakemake file (the *Snakefile)* that runs the analysis. The output of running the pipeline is placed in a results subdirectory, although only key results files are tracked in the GitHub repository since many of them are very large. The pipeline also generates HTML rendering of the key analysis notebooks and result plots, which are available at https://dms-vep.github.io/SARS-CoV-2_Omicron_BA.1_spike_DMS_mAbs_for_BA.1 and https://dms-vep.github.io/SARS-CoV-2_Delta_spike_DMS_REGN10933 for Delta. Looking at these websites is the easiest way to understand the analysis. Note that many of the plots are interactive charts created with Altair (VanderPlas et al., 2018), and we encourage readers to use the interactive features to better explore the data.

### Analysis of PacBio data to link barcodes to spike mutations

To link each barcode to its spike variant, we used *alignparse* (Crawford and Bloom, 2019) to process the PacBio CCSs to determine the barcode and spike mutations for each CCS.

We performed several quality control steps. First we examined the synonymous tags introduced at the end of each amplicon during the library preparation to identify CCSs with discordant tags indicative of strand exchange during library preparation: typically <2% of CCSs had discordant tags, indicating a low rate of strand exchange. CCSs with identified strand exchange or out of frame indels were discarded. Next, we computed the empirical accuracy of the CCSs by examining how often CCSs with the same barcode reported the same spike sequence (the exact method used to compute the empirical accuracy is implemented here: https://jbloomlab.github.io/alignparse/alignparse.consensus.html#alignparse.consensus.empirical_accuracy). The empirical accuracies were between 0.65 and 0.75, indicating that fraction of CCSs correctly report the actual mutations. The inaccuracies are due to a combination of sequencing errors, reverse transcription errors, PCR strand exchange, and occasional actual association of the same barcode with different variants in different cells (which can especially arise if the complexity of the initial virus library integrated into cells at single copy is not much higher than the complexity of the final cell library).

We then built consensus sequences for each barcode with at least three CCSs, using the method implemented at https://jbloomlab.github.io/alignparse/alignparse.consensus.html#alignparse.consensus.simple_mutconsens_us with *max_minor_sub_frac* and *max_minor_indel_frac* both set to 0.2. This approach of requiring multiple concordant CCSs to call a consensus is expected to lead to higher accuracy in the final barcode / spike variant linking, and will generally discard barcodes that are not uniquely linked to a single spike variant.

Files containing the final barcode / variant lookup tables and the analysis notebooks with resulting quality control plots are linked to the main HTML pages in the documentation for the BA.1 and Delta experiments as provided in the *Processed data* section below.

### Analysis of Illumina data to count barcodes for each variant in each experiment

For each experiment, we processed the Illumina barcode sequencing with the parser implemented at https://jbloomlab.github.io/dms_variants/dms_variants.illuminabarcodeparser.html to determine the counts of each variant in each condition. Barcoded variants were only retained for subsequent analysis if their “preselection” counts (no-antibody selection for antibody escape experiments, VSV-G-pseudotyped infections for functional selections) met some minimal count threshold specified in the *config.yaml* file of the GitHub repos for the BA.1 and Delta spikes. This thresholding removes variants that are expected to have substantial noise due to low counts. Note that a caveat that should be kept in mind is that the actual key bottleneck is expected to usually occur at the stage of infection with the virus library rather than sequencing, since the barcodes are generally sequenced to a depth that greatly exceeds the complexity of the libraries used for the infections. Therefore, although variants with low counts are expected to have more noise, the counts do not enable a quantitative estimate of the actual bottleneck size experienced by each variant.

Files containing the barcode counts and the analysis notebook with resulting quality control plots are linked to the main HTML pages in the documentation for the BA.1 and Delta experiments as provided in the *Processed data* section below.

### Computing functional effects of mutations in deep mutational scanning

To estimate the functional effects of individual mutations, we first computed functional scores for each variant from the counts in the VSV-G-pseudotyped library infection (which should not impose any selection on the spike) versus the spike-pseudotyped library infected into ACE2 expressing target cells. The functional score for variant *v* is defined as *log_2_* ([n_v__post_ / *n^wt^_post_*]/[*n^v^_pre_* / *n^wt^_pre_*]) where *n^v^_post_* is the count of variant *v* in the post-selection (spike-pseudotyped) infection, *n^v^_pre_* is the count of variant *v* in the pre-selection (VSV-G-pseudotyped) infection, and *n^wt^_post_* and *n^wt^_post_* are the counts of all unmutated (wildtype) variants in each condition. Negative functional scores indicate a spike variant is worse at mediating infection than the unmutated spike, and positive functional scores indicate a variant is better at mediating infection than the unmutated spike. The distributions of these functional scores are plotted in Figure 2.

To deconvolve the functional scores for all spike variants (which often contain multiple mutations) into estimates of the effects of individual amino-acid mutations, we fit global epistasis models (Otwinowski et al., 2018) to the variant functional scores, using the models implemented at https://jbloomlab.github.io/dms_variants/dms_variants.globalepistasis.html with the Gaussian likelihood function. This fitting estimates how each mutation affects an underlying latent phenotype, as well as the shape of the global epistasis function relating the latent phenotype to the observed functional score. We also then re-transform the inferred effect of each mutation on this latent phenotype through the global epistasis function to estimate its effect on the observed phenotype. This approach provides a way to deconvolve the information in the multi-mutant variants to more accurately estimate the effects of mutations under the assumptions of a global epistasis model (Otwinowski et al., 2018). For final reporting, we then took the average (median) of the estimated functional effect of each mutation across all the replicates and libraries for each different spike (BA.1 or Delta). In Figure 7 of this paper we report the functional effects on the observed (rather than latent) phenotype, as that is a more relevant measure of its expected impact on spike-mediated infection.

Files containing the effects of mutations on both the latent and observed phenotypes for both individual replicates/libraries and averages across them, the analysis notebooks with relevant quality control plots, and interactive plots summarizing the final estimates are linked to the main HTML pages in the documentation for the BA.1 and Delta experiments as provided in the *Processed data* section below.

For the figures in this paper and the default view of the interactive plots described in the *Processed data* section below, we only include mutations seen in at least three distinct variants (averaged across libraries).

### Computing antibody escape by mutations

For the antibody selections, we computed the non-neutralized fraction (probability of escape) *p_v_*(c) for each variant *v* at each antibody concentration *c* for a given antibody as *p_v_(c)* = *(n_v_^c^/S^c^)/(n_v_^0^/S^0^)* where *n_v_^c^* is the count of variant *v* at antibody concentration *c*, *n_v_^0^* is the count of variant *v* in the no-antibody control, S^c^ is the total counts of the neutralization standard at antibody concentration *c*, and S^0^ is the total concentration of the neutralization standard in the no-antibody control. These values should in principle fall between 0 (variant is totally neutralized by antibody) and 1 (variant is not neutralized by antibody), and in practice we clip any values measured as >1 to a value of 1.

To deconvolve mutation-level escape values from the measured *p_v_(c)* values for the variants (which often contain multiple mutations) at multiple concentrations, we used the approach implemented in *polyclonal* software package (https://jbloomlab.github.io/polyclonal/) (Yu et al., 2022), constraining the fits to a single epitope (since we are only analyzing monoclonal antibodies). This analysis yields a mutation-level escape score for each observed variant (the *β_m,e_* values in the nomenclature of the *polyclonal* package) which will be zero for mutations that have no effect on antibody escape, and >0 for mutations that mediate antibody escape. These are the values plotted in the heat maps shown in the antibody escape figures in this paper; the line plots show site-level summaries of these values (eg, the sum of the escape values at each site). Note that the *polyclonal* models (Yu et al., 2022) can use the escape values inferred from the deep mutational scanning to predict the non-neutralized fraction for arbitrary mutants, and we correlated those measurements with the IC50 values measured by standard neutralization assays in the antibody escape figures in this paper. For all antibodies we had replicate measurements (multiple libraries, and in some cases technical replicates of the same library), and the final reported values are the average (median) across these replicates.

Files containing the escape values for both individual replicates/libraries and averages across them, the analysis notebooks with relevant quality control plots, and interactive plots summarizing the final estimates are linked to the main HTML pages in the documentation for the BA.1 and Delta experiments as provided in the *Processed data* section below.

For the figures in this paper and the default view of the interactive plots described in the *Processed data* section below, we only include mutations seen in at least three distinct variants (averaged across libraries), and with a functional effect of at least −1.38 (for Omicron BA.1) or at least −1.47 (for Delta).

### Comparison of deep mutational scanning data to enrichment of mutations during actual human SARS-CoV-2 evolution

In Figure 7, we compare the functional effects of mutations measured in deep mutational scanning to the enrichment of mutations in actual human SARS-CoV-2 evolution compared that expected given the underlying viral mutation rate. The computation of the “expected” versus actual observed mutation counts is performed by the code at https://github.com/jbloomlab/SARS2-mut-rates/tree/25204ea0c868c4c86d0cc16bd5f203b1_b0607868

The approach for calculating the enrichment of observed versus expected mutations is inspired by the strategy used in (Neher, 2022). Briefly, we downloaded the pre-built UShER tree of all publicly available SARS-CoV-2 sequences (Turakhia et al., 2021) as of Sept-26-2022 from http://hgdownload.soe.ucsc.edu/goldenPath/wuhCor1/UShER_SARS-CoV-2/2022/09/26/public-2022-09-26.all.masked.nextclade.pangolin.pb.gz. We then used matUtils (Turakhia et al., 2021) to separate the tree by the pre-annotated Nextstrain clades, and extract the number of unique ***occurrences*** of each nucleotide mutation along the tree branches for each clade. Note that we are counting unique occurrences of each mutation along the tree (number of branches with a mutation), not the number of observations of the mutation in the final set of aligned mutations. This is an important distinction, as a mutation could be common in the final aligned sequences even if it only had one or a few occurrences. In counting these occurrences, we ignored mutations on any branch with more than four nucleotide mutations, or more than one nucleotide mutation to either the Wuhan-Hu-1 reference sequence or the founder sequence for that clade, as taken from (Neher, 2022). The reason for excluding mutations on branches with high numbers of mutations as these can often indicate problematic sequences; the reason for excluding mutations on branches with multiple reversions to reference is that sometimes sequences with missing coverage have identities automatically called to reference. After tabulating these mutation counts, we ignored any mutations that are reversions from the clade founder to the Wuhan-Hu-1 reference or reversions of these, a set of sites known to be subject to sequencing errors (Turakhia et al., 2020), or were at sites with unusually high mutation rates (see sites manually specified in the config.yaml file at the GitHub repo listed above). Overall, this process provided a list of the counts of nucleotide mutations for each Nextstrain clade after filtering branches and mutations/sites expected to be affected by various possible errors.

We then calculated the relative rates of each type of the 12 possible nucleotide mutations at fourfold synonymous third codon positions by counting the mutation at such positions in each Nextstrain clade, and then normalizing the fraction of these positions that had each nucleotide identity at each site, retaining only Nextstrain clades with at least 5000 such mutation counts. We then calculated for each clade the expected number of counts of each nucleotide mutation based on these rates and the actual counts at the four-fold synonymous sites. The result of these calculations is how many times we expect to observe each nucleotide mutation if all sites are under the same constraint as the four-fold synonymous sites. Next, we aggregated the expected and observed counts across all Nextstrain clades, and also across all nucleotide mutations that encoded for the same amino-acid mutation to get expected and observed counts for aminoacid mutations, and only retained amino-acid mutations with at least 20 expected counts. Figure 7D plots the log base 2 of the expected versus observed counts after adding a pseudocount of 0.5.

We compared these counts to deep mutational scanning data from our study and several other spike deep mutational scanning studies as described in the main text. The tabulated deep mutational scanning measurements and the enrichments of mutations in actual SARS-CoV-2 evolution are at https://github.com/dms-vep/SARS-CoV-2_Omicron_BA.1_spike_DMS_mAbs/blob/main/results/compare_muteffects/natural_enrichment_vs_dms.csv

### Processed data

The key results from the analysis are stored in the results subdirectory of the GitHub repos for BA.1 (https://github.com/dms-vep/SARS-CoV-2_Omicron_BA.1_spike_DMS_mAbs) and Delta (https://github.com/dms-vep/SARS-CoV-2_Delta_spike_DMS_REGN10933). The easiest way to navigate these results are via the HTML documentation at https://dms-vep.github.io/SARS-CoV-2_Omicron_BA.1_spike_DMS_mAbs and https://dms-vep.github.io/SARS-CoV-2_Delta_spike_DMS_REGN10933 These pages contain links to the key data files, as well as interactive heat maps of the functional effects of mutations and the effects of mutations on antibody escape. Note that these plots are interactive, and allow you to filter by certain regions of the protein, the number of variants in which a mutation is seen, the maximum magnitude of an effect at a given site, and other relevant parameters.

In the final output files, mutations are numbered in reference-based (Wuhan-Hu-1) spike numbering. The GitHub repos contain files that convert sequential numbering of the BA.1 and Delta spike to reference based numbering.

### Raw sequencing data

The raw PacBio and Illumina sequencing data have been deposited on the NCBI’s Sequence Read Archive with BioProject number PRJNA888402 for the Omicron BA.1 data and PRJNA889020 for the Delta data. The PacBio sequencing linking variants to barcodes can be found under BioSample accessions SAMN31220980 for Omicron BA.1 and SAMN31230634 for Delta. The Illumina barcode sequencing can be found under BioSample accessions SAMN31216920 for Omicron BA.1 and SAMN31230628 for Delta.

## Acknowledgments

We thank Michael Emerman for scientific advice and Kenneth A Matreyek for providing HEK-293T-ACE2-medium cell line. This work was supported in part by the NIH/NIAID under grant R01AI141707 and contract 75N93021C00015 to JDB. This work was supported in part by a Gates Foundation grant INV-004949 to JDB. JDB is an Investigator of the Howard Hughes Medical Institute. KHDC was supported by NIH/NIAID grant F30AI149928. BD was funded in part by EMBO under grant ALTF 81-2020. CER was supported by the Viral Pathogenesis and Evolution training grant T32 AI083203. TCY was supported by the CMB training grant T32 GM007270 and the NSF graduate research fellowship DGE-2140004. DRB was supported by NIAID grant UM1 AI44462 and Bill and Melinda Gates Foundation grants OPP1170236 and INV-004923.

---

**Supplementary Figure 1.**
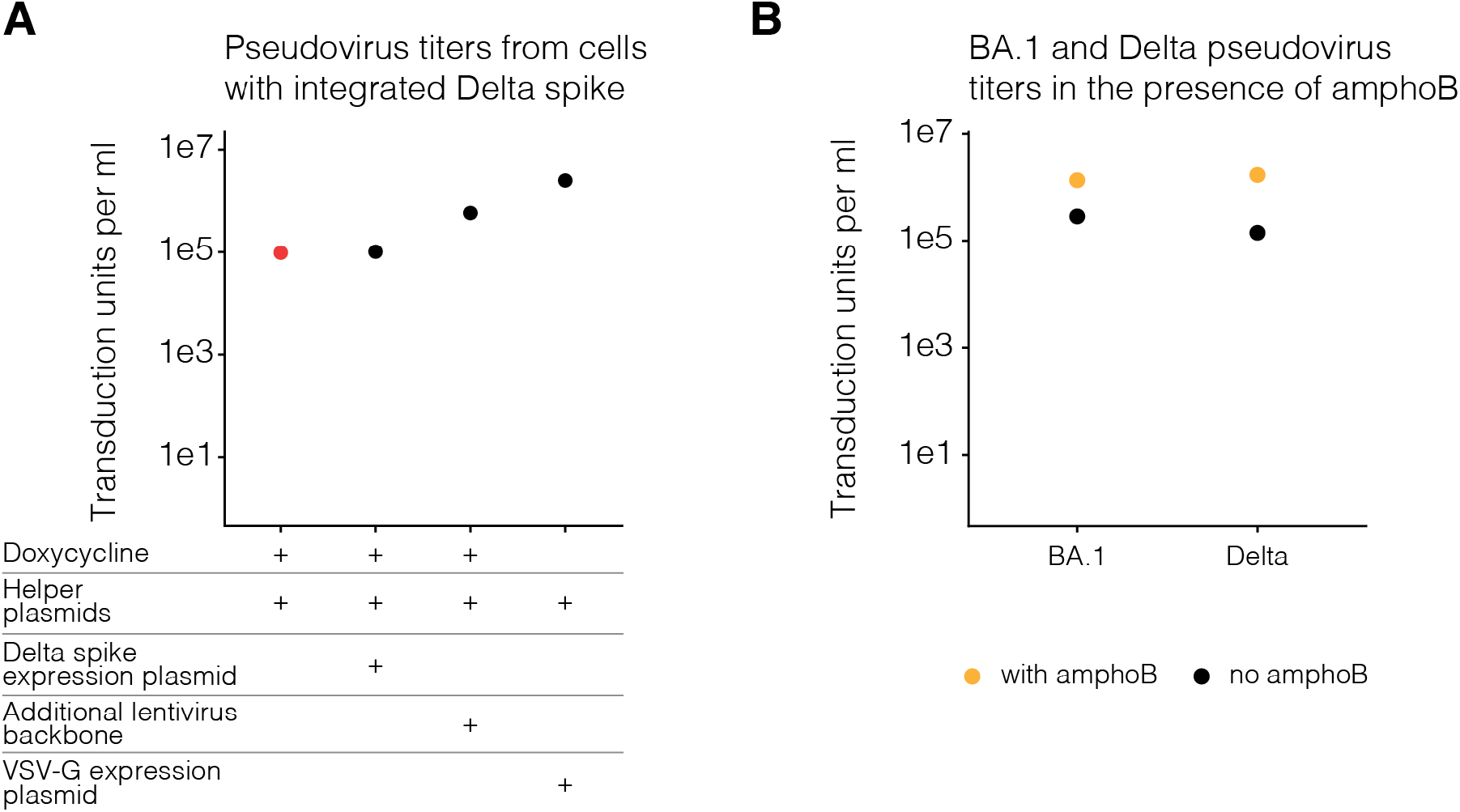
Pseudovirus titers from phenotype-genotype linked lentiviruses. **(A)** Delta spike pseudotyped lentivirus titers. Viruses were produced under indicated conditions from cells with integrated lentivirus genomes carrying Delta spike. Virus titers for conditions used to generate the actual deep mutational scanning libraries are coloured red. Viruses were titrated on ACE2-TMPRSS2-HEK-293T cells. **(B)** BA.1 or Delta spike-pseudotyped lentivirus titers in the presence or absence of amphotericin B (amphoB). BA.1 virus was titrated on ACE2-HEK-293T cells and Delta virus was titrated on ACE2-TMPRSS2-HEK-293T cells.

**Supplementary Figure 2.**
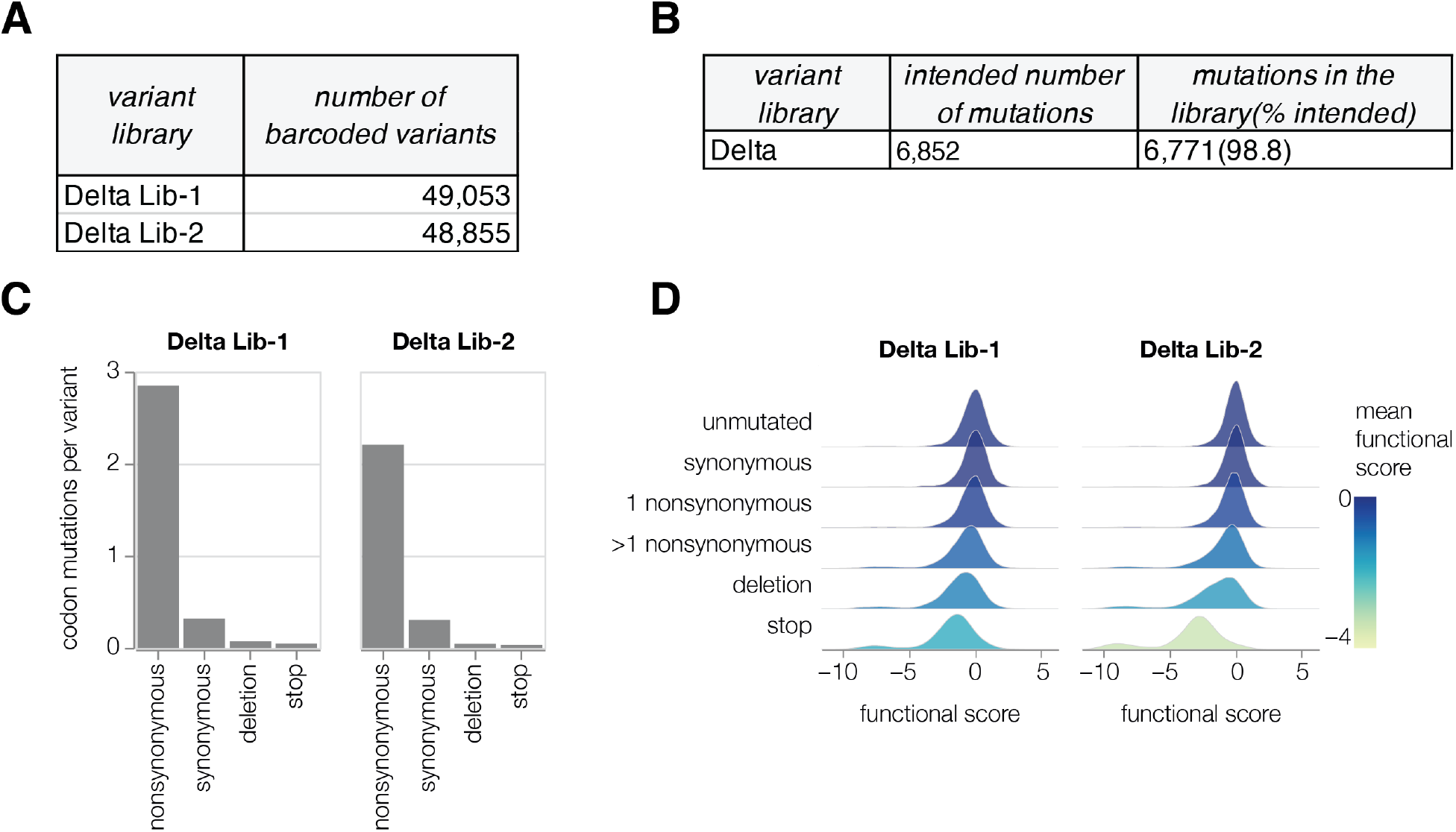
Delta spike deep mutational scanning libraries. **(A)** Total number of barcoded variants in each Delta library. **(B)** Coverage of intended mutations across both Delta libraries. **(C)** Average number of mutations per barcoded spike in Delta libraries. **(D)** Distribution of functional scores for variants with different types of mutations in the Delta libraries.

**Supplementary Figure 3.**
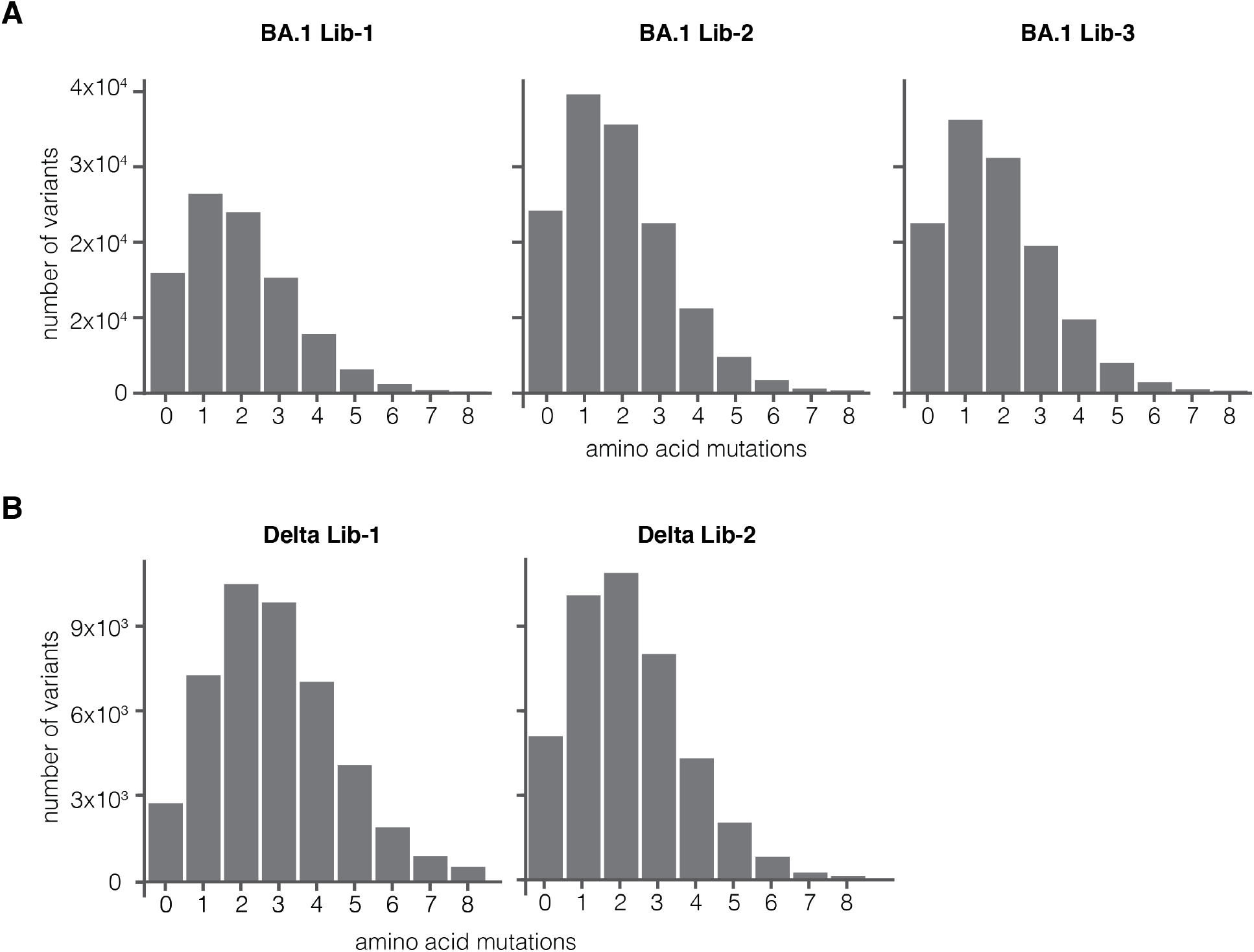
Distribution of the number of amino-acid mutations per variant in BA.1 (A) and Delta (B) deep mutational scanning libraries.

**Supplementary Figure 4.**
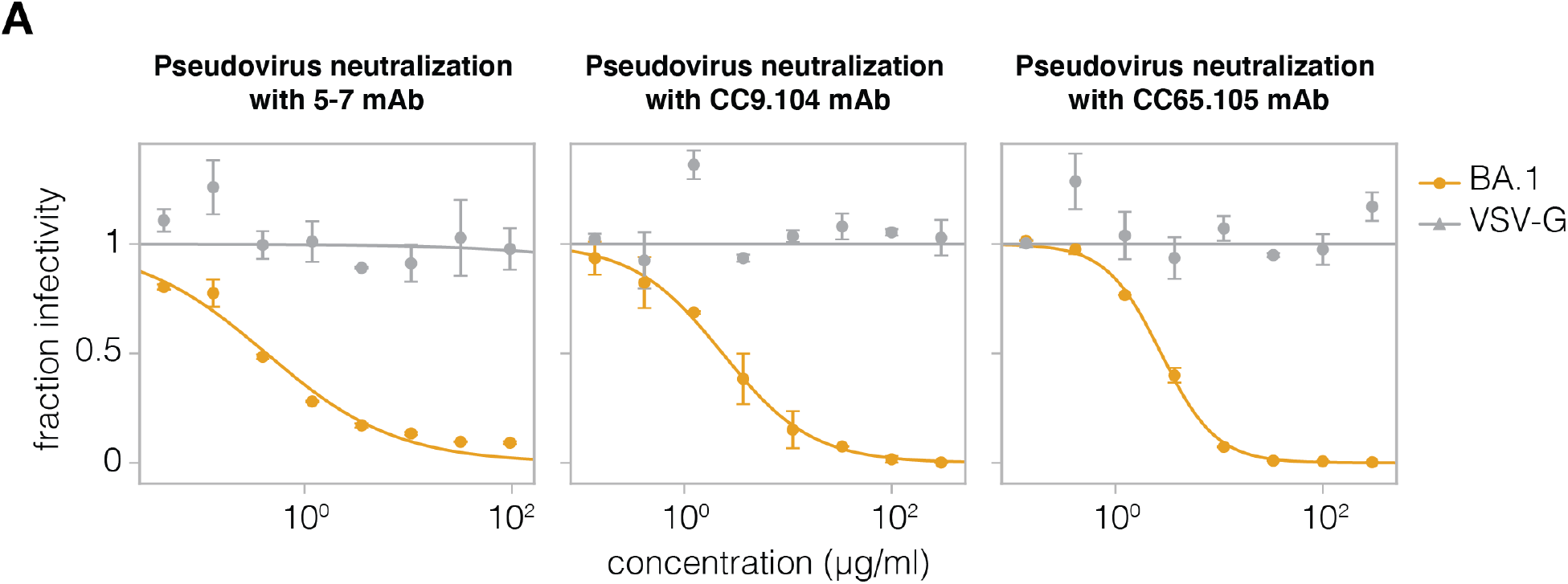
The VSV-G neutralization standard is not neutralized by antibodies 5-7, CC9.104 and CC65.105. **(A)** Neutralization assays using NTD-targeting 5-7 mAb and S2-targeting CC9.104 and CC65.105 antibodies against lentivirus pseudotyped with BA.1 spike or VSV-G.

**Supplementary Figure 5.**
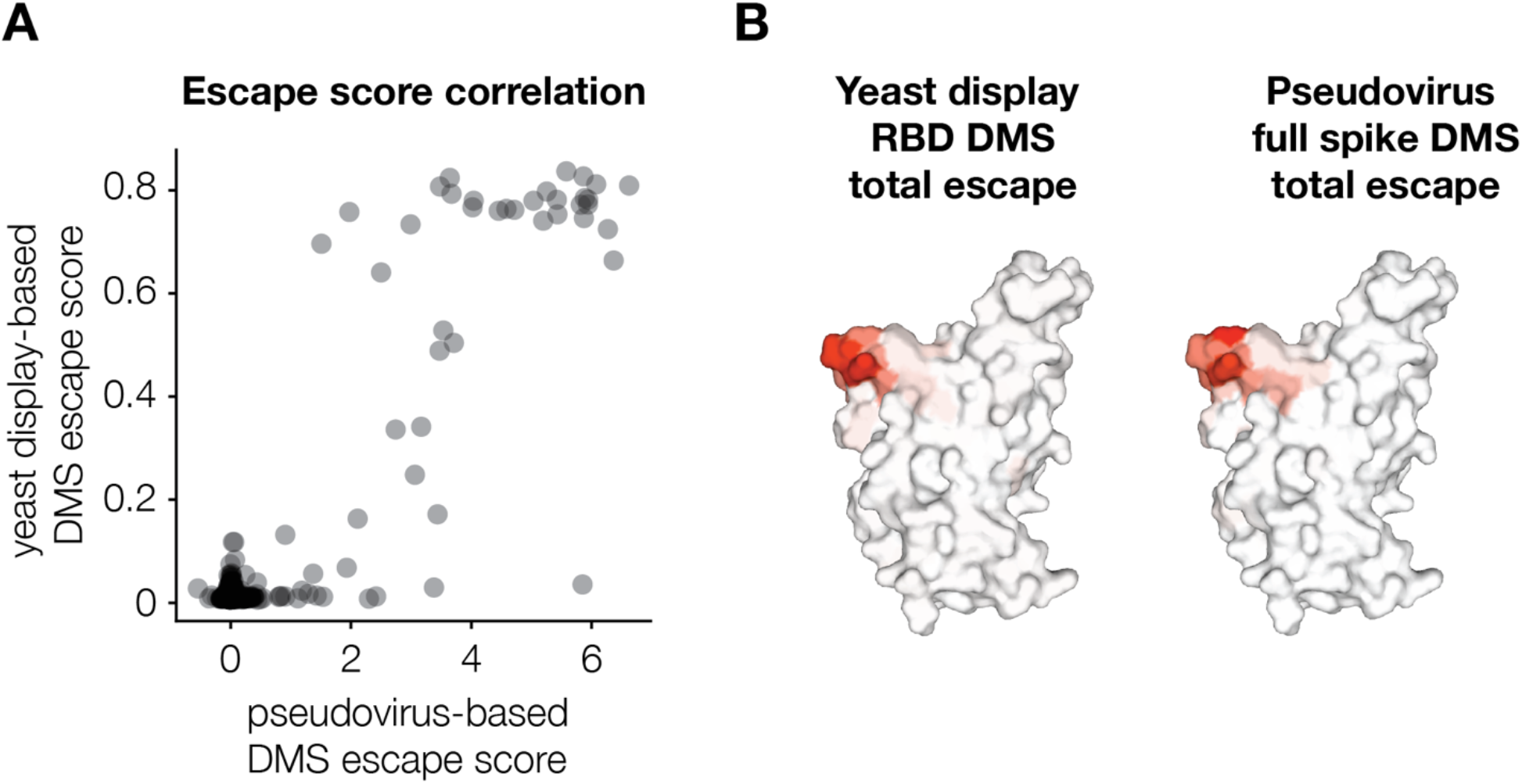
Comparison between LY-CoV1404 escape mapping using full spike pseudovirus deep mutational scanning versus our previously described yeastdisplay deep mutational scanning of just the RBD. **(A)** Correlation between measured mutation-level escape scores for LY-CoV1404 antibody in pseudovirus and yeast display deep mutational scanning experiments. Yeast display data is taken from (Starr et al., 2022). **(B)** Surface representation of SARS-CoV-2 RBD coloured by sum of escape scores at that site. PDB ID: 6XM4.

**Supplementary Figure 6.**
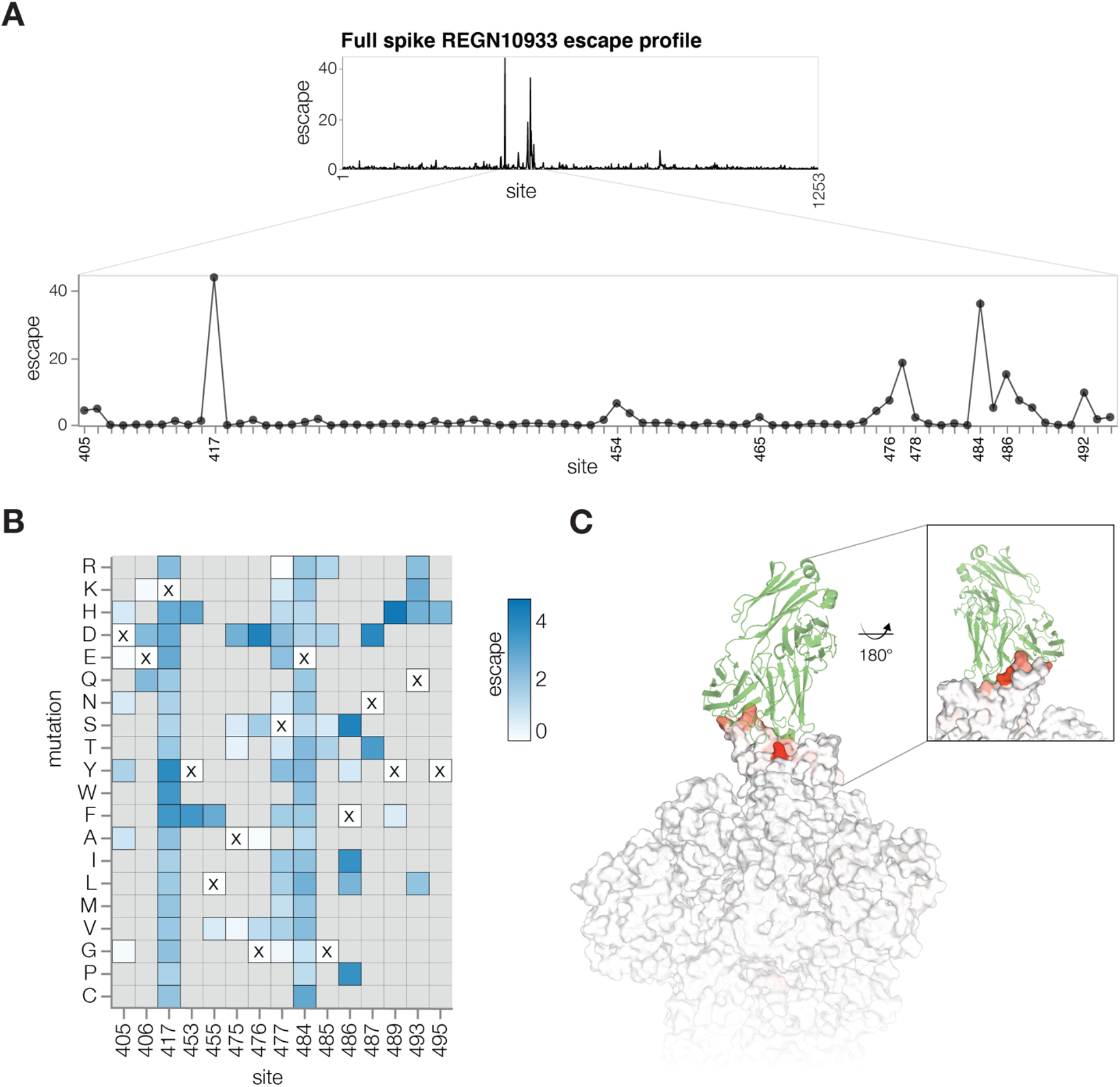
Antibody REGN10933 escape mapping using Delta deep mutational scanning libraries. **(A)** Total escape scores for each site within Delta spike and a zoomed-in plot showing key escape sites. **(B)** Heatmap of mutation escape scores at key sites. Residues marked with X are the wildtype amino acids in Delta sequence. Amino acids not present in our libraries are shown in gray. An interactive version of this heatmap for the entirety of spike is at https://dms-vep.github.io/SARS-CoV-2_Delta_spike_DMS_REGN10933/REGN10933_escape_plot.html **(C)** Surface representation of spike coloured by sum of escape scores at that site. REGN10933 antibody is shown in green. PDB structures 6XDG and 6XM4 were aligned to make this figure. Site numbering in all plots is based on the Wuhan-Hu-1 sequence.

**Supplementary Table 1.**
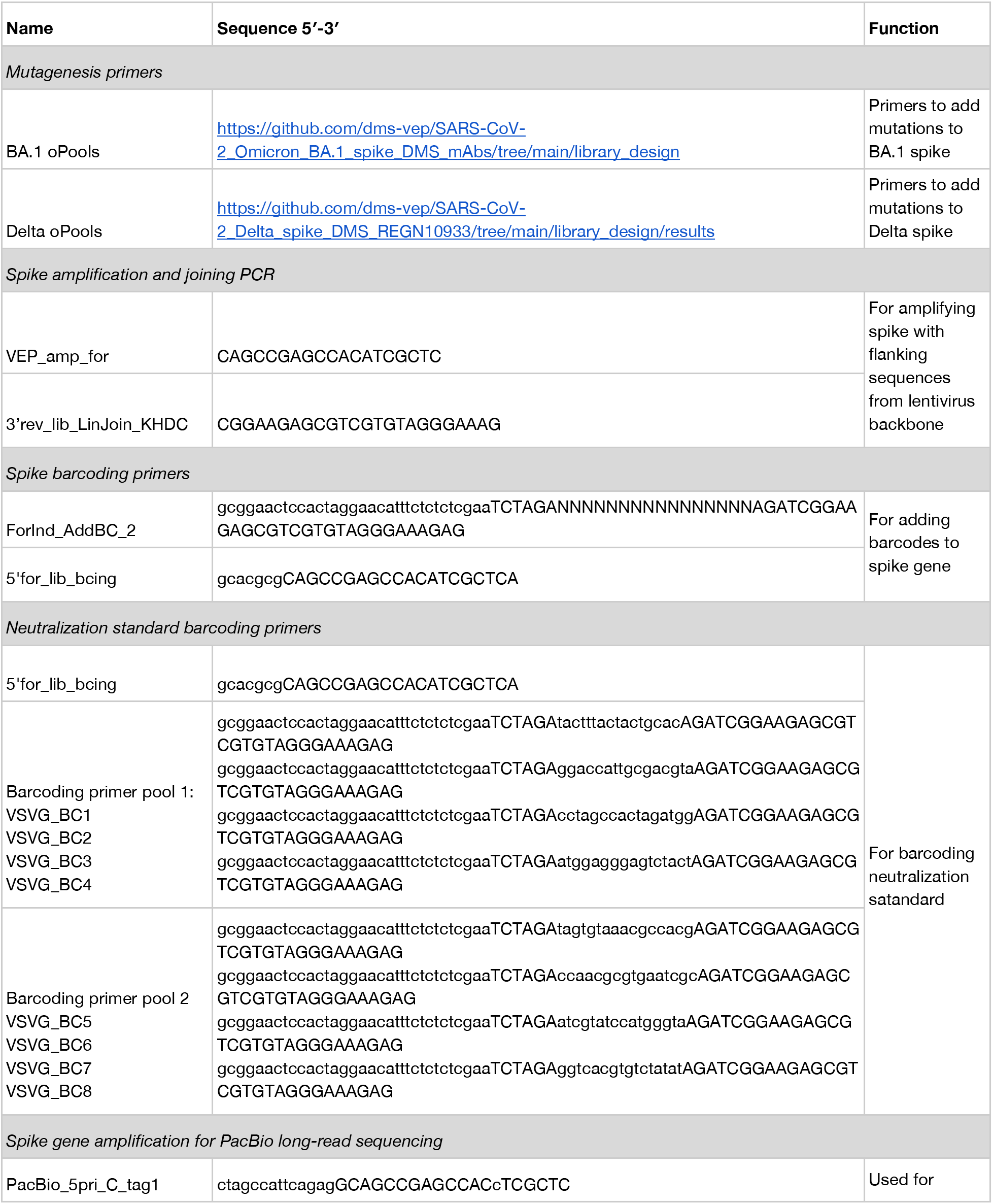
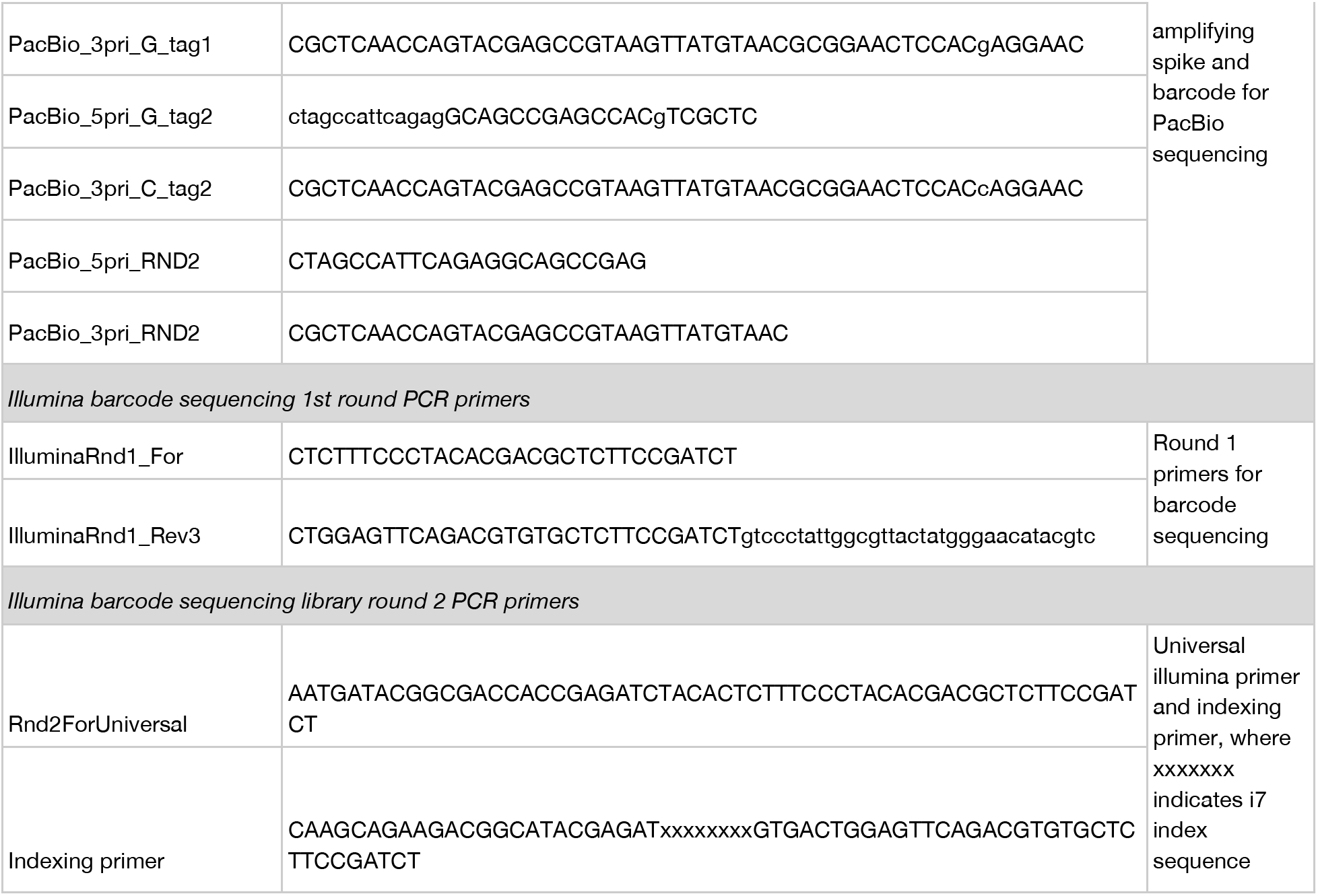
Primer sequences used for building deep mutational scanning libraries.

## References

Ballal, A., Malliaris, C.D., Morozov, A.V., 2020. Molecular Phenotypes as Key Intermediates in Mapping Genotypes to Fitness, in: Pontarotti, P. (Ed.), Evolutionary Biology—A Transdisciplinary Approach. Springer International Publishing, Cham, pp. 15–40. https://doi.org/10.1007/978-3-030-57246-4_2

Baum, A., Fulton, B.O., Wloga, E., Copin, R., Pascal, K.E., Russo, V., Giordano, S., Lanza, K., Negron, N., Ni, M., Wei, Y., Atwal, G.S., Murphy, A.J., Stahl, N., Yancopoulos, G.D., Kyratsous, C.A., 2020. Antibody cocktail to SARS-CoV-2 spike protein prevents rapid mutational escape seen with individual antibodies. Science eabd0831. https://doi.org/10.1126/science.abd0831

Bellusci, L., Grubbs, G., Zahra, F.T., Forgacs, D., Golding, H., Ross, T.M., Khurana, S., 2022. Antibody affinity and cross-variant neutralization of SARS-CoV-2 Omicron BA.1, BA.2 and BA.3 following third mRNA vaccination. Nat. Commun. 13, 4617. https://doi.org/10.1038/s41467-022-32298-w

Benton, D.J., Wrobel, A.G., Roustan, C., Borg, A., Xu, P., Martin, S.R., Rosenthal, P.B., Skehel, J.J., Gamblin, S.J., 2021. The effect of the D614G substitution on the structure of the spike glycoprotein of SARS-CoV-2. Proc. Natl. Acad. Sci. 118, e2022586118. https://doi.org/10.1073/pnas.2022586118

Bloom, J.D., 2014. An Experimentally Determined Evolutionary Model Dramatically Improves Phylogenetic Fit. Mol. Biol. Evol. 31, 1956–1978. https://doi.org/10.1093/molbev/msu173

Cao, Y., Jian, F., Wang, J., Yu, Y., Song, W., Yisimayi, A., Wang, J., An, R., Zhang, N., Wang, Yao, Wang, P., Zhao, L., Sun, H., Yu, L., Yang, S., Niu, X., Xiao, T., Gu, Q., Shao, F., Hao, X., Xu, Y., Jin, R., Wang, Youchun, Xie, X.S., 2022. Imprinted SARS-CoV-2 humoral immunity induces converging Omicron RBD evolution. https://doi.org/10.1101/2022.09.15.507787

Cerutti, G., Guo, Y., Wang, P., Nair, M.S., Wang, M., Huang, Y., Yu, J., Liu, L., Katsamba, P.S., Bahna, F., Reddem, E.R., Kwong, P.D., Ho, D.D., Sheng, Z., Shapiro, L., 2021. Neutralizing antibody 5-7 defines a distinct site of vulnerability in SARS-CoV-2 spike N-terminal domain. Cell Rep. 37, 109928. https://doi.org/10.1016/j.celrep.2021.109928

Chen, C., Nadeau, S., Yared, M., Voinov, P., Xie, N., Roemer, C., Stadler, T., 2022. CoV-Spectrum: analysis of globally shared SARS-CoV-2 data to identify and characterize new variants. Bioinformatics 38, 1735–1737. https://doi.org/10.1093/bioinformatics/btab856

Chun, T.-W., Carruth, L., Finzi, D., Shen, X., DiGiuseppe, J.A., Taylor, H., Hermankova, M., Chadwick, K., Margolick, J., Quinn, T.C., Kuo, Y.-H., Brookmeyer, R., Zeiger, M.A., Barditch-Crovo, P., Siliciano, R.F., 1997. Quantification of latent tissue reservoirs and total body viral load in HIV-1 infection. Nature 387, 183–188. https://doi.org/10.1038/387183a0

Crawford, K.H.D., Bloom, J.D., 2019. alignparse: A Python package for parsing complex features from high-throughput long-read sequencing. J. Open Source Softw. 4, 1915. https://doi.org/10.21105/joss.01915

Crawford, K.H.D., Dingens, A.S., Eguia, R., Wolf, C.R., Wilcox, N., Logue, J.K., Shuey, K., Casto, A.M., Fiala, B., Wrenn, S., Pettie, D., King, N.P., Greninger, A.L., Chu, H.Y., Bloom, J.D., 2021. Dynamics of Neutralizing Antibody Titers in the Months After Severe Acute Respiratory Syndrome Coronavirus 2 Infection. J. Infect. Dis. 223, 197–205. https://doi.org/10.1093/infdis/jiaa618

Crawford, K.H.D., Eguia, R., Dingens, A.S., Loes, A.N., Malone, K.D., Wolf, C.R., Chu, H.Y., Tortorici, M.A., Veesler, D., Murphy, M., Pettie, D., King, N.P., Balazs, A.B., Bloom, J.D., 2020. Protocol and Reagents for Pseudotyping Lentiviral Particles with SARS-CoV-2 Spike Protein for Neutralization Assays. Viruses 12, 513. https://doi.org/10.3390/v12050513

Cronin, J., Zhang, X.-Y., Reiser, J., 2005. Altering the Tropism of Lentiviral Vectors through Pseudotyping. Curr. Gene Ther. 5, 387–398.

DeGrace, M.M., Ghedin, E., Frieman, M.B., Krammer, F., Grifoni, A., Alisoltani, A., Alter, G., Amara, R.R., Baric, R.S., Barouch, D.H., Bloom, J.D., Bloyet, L.-M., Bonenfant, G., Boon, A.C.M., Boritz, E.A., Bratt, D.L., Bricker, T.L., Brown, L., Buchser, W.J., Carreño, J.M., Cohen-Lavi, L., Darling, T.L., Davis-Gardner, M.E., Dearlove, B.L., Di, H., Dittmann, M., Doria-Rose, N.A., Douek, D.C., Drosten, C., Edara, V.-V., Ellebedy, A., Fabrizio, T.P., Ferrari, G., Fischer, W.M., Florence, W.C., Fouchier, R. A.M., Franks, J., García-Sastre, A., Godzik, A., Gonzalez-Reiche, A.S., Gordon, A., Haagmans, B.L., Halfmann, P.J., Ho, D.D., Holbrook, M.R., Huang, Y., James, S.L., Jaroszewski, L., Jeevan, T., Johnson, R.M., Jones, T.C., Joshi, A., Kawaoka, Y., Kercher, L., Koopmans, M.P.G., Korber, B., Koren, E., Koup, R.A., LeGresley, E.B., Lemieux, J.E., Liebeskind, M.J., Liu, Z., Livingston, B., Logue, J.P., Luo, Y., McDermott, A.B., McElrath, M.J., Meliopoulos, V.A., Menachery, V.D., Montefiori, D.C., Mühlemann, B., Munster, V.J., Munt, J.E., Nair, M.S., Netzl, A., Niewiadomska, A.M., O’Dell, S., Pekosz, A., Perlman, S., Pontelli, M.C., Rockx, B., Rolland, M., Rothlauf, P.W., Sacharen, S., Scheuermann, R.H., Schmidt, S.D., Schotsaert, M., Schultz-Cherry, S., Seder, R.A., Sedova, M., Sette, A., Shabman, R.S., Shen, X., Shi, P.-Y., Shukla, M., Simon, V., Stumpf, S., Sullivan, N.J., Thackray, L.B., Theiler, J., Thomas, P.G., Trifkovic, S., Türeli, S., Turner, S.A., Vakaki, M.A., van Bakel, H., VanBlargan, L.A., Vincent, L.R., Wallace, Z.S., Wang, L., Wang, M., Wang, P., Wang, W., Weaver, S.C., Webby, R.J., Weiss, C.D., Wentworth, D.E., Weston, S.M., Whelan, S.P.J., Whitener, B.M., Wilks, S.H., Xie, X., Ying, B., Yoon, H., Zhou, B., Hertz, T., Smith, D.J., Diamond, M.S., Post, D.J., Suthar, M.S., 2022. Defining the risk of SARS-CoV-2 variants on immune protection. Nature 605, 640–652. https://doi.org/10.1038/s41586-022-04690-5

Dingens, A.S., Acharya, P., Haddox, H.K., Rawi, R., Xu, K., Chuang, G.-Y., Wei, H., Zhang, B., Mascola, J.R., Carragher, B., Potter, C.S., Overbaugh, J., Kwong, P.D., Bloom, J.D., 2018. Complete functional mapping of infection- and vaccine-elicited antibodies against the fusion peptide of HIV. PLOS Pathog. 14, e1007159. https://doi.org/10.1371/iournal.ppat.1007159

Farrell, A.G., Dadonaite, B., Greaney, A.J., Eguia, R., Loes, A.N., Franko, N.M., Logue, J., Carreño, J.M., Abbad, A., Chu, H.Y., Matreyek, K.A., Bloom, J.D., 2022. Receptor-Binding Domain (RBD) Antibodies Contribute More to SARS-CoV-2 Neutralization When Target Cells Express High Levels of ACE2. Viruses 14, 2061. https://doi.org/10.3390/v14092061

Feng, S., Phillips, D.J., White, T., Sayal, H., Aley, P.K., Bibi, S., Dold, C., Fuskova, M., Gilbert, S.C., Hirsch, I., Humphries, H.E., Jepson, B., Kelly, E.J., Plested, E., Shoemaker, K., Thomas, K.M., Vekemans, J., Villafana, T.L., Lambe, T., Pollard, A.J., Voysey, M., 2021. Correlates of protection against symptomatic and asymptomatic SARS-CoV-2 infection. Nat. Med. 27, 2032–2040. https://doi.org/10.1038/s41591-021-01540-1

Gilbert, P.B., Montefiori, D.C., McDermott, A.B., Fong, Y., Benkeser, D., Deng, W., Zhou, H., Houchens, C.R., Martins, K., Jayashankar, L., Castellino, F., Flach, B., Lin, B.C., O’Connell, S., McDanal, C., Eaton, A., Sarzotti-Kelsoe, M., Lu, Y., Yu, C., Borate, B., van der Laan, L.W.P., Hejazi, N.S., Huynh, C., Miller, J., El Sahly, H.M., Baden, L.R., Baron, M., De La Cruz, L., Gay, C., Kalams, S., Kelley, C.F., Andrasik, M.P., Kublin, J.G., Corey, L., Neuzil, K.M., Carpp, L.N., Pajon, R., Follmann, D., Donis, R.O., Koup, R.A., 2022. Immune correlates analysis of the mRNA-1273 COVID-19 vaccine efficacy clinical trial. Science 375, 43–50. https://doi.org/10.1126/science.abm3425

Greaney, A.J., Starr, T.N., Barnes, C.O., Weisblum, Y., Schmidt, F., Caskey, M., Gaebler, C., Cho, A., Agudelo, M., Finkin, S., Wang, Z., Poston, D., Muecksch, F., Hatziioannou, T., Bieniasz, P.D., Robbiani, D.F., Nussenzweig, M.C., Bjorkman, P.J., Bloom, J.D., 2021. Mapping mutations to the SARS-CoV-2 RBD that escape binding by different classes of antibodies. Nat. Commun. 12, 4196. https://doi.org/10.1038/s41467-021-24435-8

Greaney, A.J., Starr, T.N., Bloom, J.D., 2022. An antibody-escape estimator for mutations to the SARS-CoV-2 receptor-binding domain. Virus Evol. 8, veac021. https://doi.org/10.1093/ve/veac021

Haddox, H.K., Dingens, A.S., Bloom, J.D., 2016. Experimental Estimation of the Effects of All Amino-Acid Mutations to HIV’s Envelope Protein on Viral Replication in Cell Culture. PLOS Pathog. 12, e1006114. https://doi.org/10.1371/journal.ppat.1006114

Hansen, J., Baum, A., Pascal, K.E., Russo, V., Giordano, S., Wloga, E., Fulton, B.O., Yan, Y., Koon, K., Patel, K., Chung, K.M., Hermann, A., Ullman, E., Cruz, J., Rafique, A., Huang, T., Fairhurst, J., Libertiny, C., Malbec, M., Lee, W., Welsh, R., Farr, G., Pennington, S., Deshpande, D., Cheng, J., Watty, A., Bouffard, P., Babb, R., Levenkova, N., Chen, C., Zhang, B., Romero Hernandez, A., Saotome, K., Zhou, Y., Franklin, M., Sivapalasingam, S., Lye, D.C., Weston, S., Logue, J., Haupt, R., Frieman, M., Chen, G., Olson, W., Murphy, A.J., Stahl, N., Yancopoulos, G.D., Kyratsous, C.A., 2020. Studies in humanized mice and convalescent humans yield a SARS-CoV-2 antibody cocktail. Science 369, 1010–1014. https://doi.org/10.1126/science.abd0827

Harms, M.J., Thornton, J.W., 2013. Evolutionary biochemistry: revealing the historical and physical causes of protein properties. Nat. Rev. Genet. 14, 559–571. https://doi.org/10.1038/nrg3540

Havranek, K.E., Jimenez, A.R., Acciani, M.D., Lay Mendoza, M.F., Reyes Ballista, J.M., Diaz, D.A., Brindley, M.A., 2020. SARS-CoV-2 Spike Alterations Enhance Pseudoparticle Titers and Replication-Competent VSV-SARS-CoV-2 Virus. Viruses 12, 1465. https://doi.org/10.3390/v12121465

Hiatt, J.B., Patwardhan, R.P., Turner, E.H., Lee, C., Shendure, J., 2010. Parallel, tag-directed assembly of locally derived short sequence reads. Nat. Methods 7, 119–122. https://doi.org/10.1038/nmeth.1416

Hill, A.J., McFaline-Figueroa, J.L., Starita, L.M., Gasperini, M.J., Matreyek, K.A., Packer, J., Jackson, D., Shendure, J., Trapnell, C., 2018. On the design of CRISPR-based single-cell molecular screens. Nat. Methods 15, 271–274. https://doi.org/10.1038/nmeth.4604

Huang, S.-W., Tai, C.-H., Hsu, Y.-M., Cheng, D., Hung, S.-J., Chai, K.M., Wang, Y.-F., Wang, J.-R., 2020. Assessing the application of a pseudovirus system for emerging SARS-CoV-2 and re-emerging avian influenza virus H5 subtypes in vaccine development. Biomed. J. 43, 375–387. https://doi.org/10.1016/j.bj.2020.06.003

Inglesby, T., Bloom, J., Brownstein, J., Burke, D., Cicero, A., Correa, D., Rio, C. del, Diggans, J., Esvelt, K., Evans, N., Evans, S.W., Fisher, M.A., Fraser, C., Goldman, E., Hoyt, K., Bloom, B.R., Lynn Klotz, Duc, J.L., Lipsitch, M., Luby, S., Morrison, S., Osterholm, M., Palmer, M.J., Pannu, J., Parthemore, C., Perkovich, G., Plotkin, S., Poste, G., Relman, D., Salzberg, S., Simonsen, L., Stearns, T., Andrew C. Weber, Yassif, J., 2022. Recommendations to Strengthen the US Government’s Enhanced Potential Pandemic Pathogen Framework and Dual Use Research of Concern Policies. https://www.centerforhealthsecurity.org/news/center-news/pdfs/220629-RecstostrengthenUSGePPPandDURCPolicies.pdf

Javanmardi, K., Chou, C.-W., Terrace, C.I., Annapareddy, A., Kaoud, T.S., Guo, Q., Lutgens, J., Zorkic, H., Horton, A.P., Gardner, E.C., Nguyen, G., Boutz, D.R., Goike, J., Voss, W.N., Kuo, H.-C., Dalby, K.N., Gollihar, J.D., Finkelstein, I.J., 2021. Rapid characterization of spike variants via mammalian cell surface display. Mol. Cell 81, 5099–5111.e8. https://doi.org/10.1016/j.molcel.2021.11.024

Jetzt, A.E., Yu, H., Klarmann, G.J., Ron, Y., Preston, B.D., Dougherty, J.P., 2000. High Rate of Recombination throughout the Human Immunodeficiency Virus Type 1 Genome. J. Virol. 74, 1234–1240.

Khare, S., Gurry, C., Freitas, L., B Schultz, M., Bach, G., Diallo, A., Akite, N., Ho, J., TC Lee, R., Yeo, W., Core Curation Team, G., Maurer-Stroh, S., GISAID Global Data Science Initiative (GISAID), Munich, Germany, Bioinformatics Institute, Agency for Science Technology and Research, Singapore, Oswaldo Cruz Foundation (FIOCRUZ), Rio de Janeiro, Brazil, Institut Pasteur de Dakar, Dakar, Senegal, National Institutes of Biotechnology Malaysia, Selangor, Malaysia, Smorodintsev Research Institute of Influenza, St. Petersburg, Russia, Genome Institute of Singapore, Agency for Science Technology and Research, Singapore, China National GeneBank, Shenzhen, China, A*STAR Infectious Disease Labs (ID Labs), Singapore, National Public Health Laboratory, National Centre for Infectious Diseases, Ministry of Health, Singapore, Department of Biological Sciences, National University of Singapore, Singapore, 2021. GISAID’s Role in Pandemic Response. China CDC Wkly. 3, 1049–1051. https://doi.org/10.46234/ccdcw2021.255

Khetawat, D., Broder, C.C., 2010. A Functional Henipavirus Envelope Glycoprotein Pseudotyped Lentivirus Assay System. Virol. J. 7, 312. https://doi.org/10.1186/1743-422X-7-312

Kobinger, G.P., Weiner, D.J., Yu, Q.-C., Wilson, J.M., 2001. Filovirus-pseudotyped lentiviral vector can efficiently and stably transduce airway epithelia in vivo. Nat. Biotechnol. 19, 225–230. https://doi.org/10.1038/85664

Larson, R.A., Dai, D., Hosack, V.T., Tan, Y., Bolken, T.C., Hruby, D.E., Amberg, S.M., 2008. Identification of a Broad-Spectrum Arenavirus Entry Inhibitor. J. Virol. 82, 10768–10775. https://doi.org/10.1128/JVI.00941-08

Lin, T.-Y., Chin, C.R., Everitt, A.R., Clare, S., Perreira, J.M., Savidis, G., Aker, A.M., John, S.P., Sarlah, D., Carreira, E.M., Elledge, S.J., Kellam, P., Brass, A.L., 2013. Amphotericin B Increases Influenza A Virus Infection by Preventing IFITM3-Mediated Restriction. Cell Rep. 5, 895–908. https://doi.org/10.1016/j.celrep.2013.10.033

Liu, J., Song, H., Liu, D., Zuo, T., Lu, F., Zhuang, H., Gao, F., 2014. Extensive Recombination Due to Heteroduplexes Generates Large Amounts of Artificial Gene Fragments during PCR. PLoS ONE 9, e106658. https://doi.org/10.1371/journal.pone.0106658

Liu, L., Wang, P., Nair, M.S., Yu, J., Rapp, M., Wang, Q., Luo, Y., Chan, J.F.-W., Sahi, V., Figueroa, A., Guo, X.V., Cerutti, G., Bimela, J., Gorman, J., Zhou, T., Chen, Z., Yuen, K.-Y., Kwong, P.D., Sodroski, J.G., Yin, M.T., Sheng, Z., Huang, Y., Shapiro, L., Ho, D.D., 2020. Potent neutralizing antibodies against multiple epitopes on SARS-CoV-2 spike. Nature 584, 450–456. https://doi.org/10.1038/s41586-020-2571-7

Liu, Lihong, Iketani, S., Guo, Y., Chan, J.F.-W., Wang, M., Liu, Liyuan, Luo, Y., Chu, H., Huang, Yiming, Nair, M.S., Yu, J., Chik, K.K.-H., Yuen, T.T.-T., Yoon, C., To, K.K.-W., Chen, H., Yin, M.T., Sobieszczyk, M.E., Huang, Yaoxing, Wang, H.H., Sheng, Z., Yuen, K.-Y., Ho, D.D., 2022. Striking antibody evasion manifested by the Omicron variant of SARS-CoV-2. Nature 602, 676–681. https://doi.org/10.1038/s41586-021-04388-0

Maher, M.C., Bartha, I., Weaver, S., di Iulio, J., Ferri, E., Soriaga, L., Lempp, F.A., Hie, B.L., Bryson, B., Berger, B., Robertson, D.L., Snell, G., Corti, D., Virgin, H.W., Kosakovsky Pond, S.L., Telenti, A., 2022. Predicting the mutational drivers of future SARS-CoV-2 variants of concern. Sci. Transl. Med. 14, eabk3445. https://doi.org/10.1126/scitranslmed.abk3445

Matreyek, K.A., Starita, L.M., Stephany, J.J., Martin, B., Chiasson, M.A., Gray, V.E., Kircher, M., Khechaduri, A., Dines, J.N., Hause, R.J., Bhatia, S., Evans, W.E., Relling, M.V., Yang, W., Shendure, J., Fowler, D.M., 2018. Multiplex Assessment of Protein Variant Abundance by Massively Parallel Sequencing. Nat. Genet. 50, 874–882. https://doi.org/10.1038/s41588-018-0122-z

McCallum, M., De Marco, A., Lempp, F.A., Tortorici, M.A., Pinto, D., Walls, A.C., Beltramello, M., Chen, A., Liu, Z., Zatta, F., Zepeda, S., di Iulio, J., Bowen, J.E., Montiel-Ruiz, M., Zhou, J., Rosen, L.E., Bianchi, S., Guarino, B., Fregni, C.S., Abdelnabi, R., Foo, S.-Y.C., Rothlauf, P.W., Bloyet, L.-M., Benigni, F., Cameroni, E., Neyts, J., Riva, A., Snell, G., Telenti, A., Whelan, S.P.J., Virgin, H.W., Corti, D., Pizzuto, M.S., Veesler, D., 2021. N-terminal domain antigenic mapping reveals a site of vulnerability for SARS-CoV-2. Cell 184, 2332–2347.e16. https://doi.org/10.1016/j.cell.2021.03.028

McCarthy, K.R., Rennick, L.J., Nambulli, S., Robinson-McCarthy, L.R., Bain, W.G., Haidar, G., Duprex, W.P., 2021. Recurrent deletions in the SARS-CoV-2 spike glycoprotein drive antibody escape. Science 371, 1139–1142. https://doi.org/10.1126/science.abf6950

Medina, M.F., Kobinger, G.P., Rux, J., Gasmi, M., Looney, D.J., Bates, P., Wilson, J.M., 2003. Lentiviral vectors pseudotyped with minimal filovirus envelopes increased gene transfer in murine lung. Mol. Ther. 8, 777–789. https://doi.org/10.1016/j.ymthe.2003.07.003

Mölder, F., Jablonski, K.P., Letcher, B., Hall, M.B., Tomkins-Tinch, C.H., Sochat, V., Forster, J., Lee, S., Twardziok, S.O., Kanitz, A., Wilm, A., Holtgrewe, M., Rahmann, S., Nahnsen, S., Köster, J., 2021. Sustainable data analysis with Snakemake. https://doi.org/10.12688/f1000research.29032.2

Naldini, L., Blömer, U., Gallay, P., Ory, D., Mulligan, R., Gage, F.H., Verma, I.M., Trono, D., 1996. In Vivo Gene Delivery and Stable Transduction of Nondividing Cells by a Lentiviral Vector. Science 272, 263–267. https://doi.org/10.1126/science.272.5259.263

Neher, R.A., 2022. Contributions of adaptation and purifying selection to SARS-CoV-2 evolution. https://doi.org/10.1101/2022.08.22.504731

OhAinle, M., Helms, L., Vermeire, J., Roesch, F., Humes, D., Basom, R., Delrow, J.J., Overbaugh, J., Emerman, M., 2018. A virus-packageable CRISPR screen identifies host factors mediating interferon inhibition of HIV. eLife 7, e39823. https://doi.org/10.7554/eLife.39823

Omelina, E.S., Ivankin, A.V., Letiagina, A.E., Pindyurin, A.V., 2019. Optimized PCR conditions minimizing the formation of chimeric DNA molecules from MPRA plasmid libraries. BMC Genomics 20, 536. https://doi.org/10.1186/s12864-019-5847-2

Otwinowski, J., McCandlish, D.M., Plotkin, J.B., 2018. Inferring the shape of global epistasis. Proc. Natl. Acad. Sci. 115, E7550–E7558. https://doi.org/10.1073/pnas.1804015115

Ouyang, W.O., Tan, T.J.C., Lei, R., Song, G., Kieffer, C., Andrabi, R., Matreyek, K.A., Wu, N.C., 2022. Probing the biophysical constraints of SARS-CoV-2 spike N-terminal domain using deep mutational scanning. https://doi.org/10.1101/2022.06.20.496903

Pang, S., Koyanagi, Y., Miles, S., Wiley, C., Vinters, H.V., Chen, I.S.Y., 1990. High levels of unintegrated HIV-1 DNA in brain tissue of AIDS dementia patients. Nature 343, 85–89. https://doi.org/10.1038/343085a0

Peacock, T.P., Goldhill, D.H., Zhou, J., Baillon, L., Frise, R., Swann, O.C., Kugathasan, R., Penn, R., Brown, J.C., Sanchez-David, R.Y., Braga, L., Williamson, M.K., Hassard, J.A., Staller, E., Hanley, B., Osborn, M., Giacca, M., Davidson, A.D., Matthews, D.A., Barclay, W.S., 2021. The furin cleavage site in the SARS-CoV-2 spike protein is required for transmission in ferrets. Nat. Microbiol. 6, 899–909. https://doi.org/10.1038/s41564-021-00908-w

Plante, J.A., Liu, Y., Liu, J., Xia, H., Johnson, B.A., Lokugamage, K.G., Zhang, X., Muruato, A.E., Zou, J., Fontes-Garfias, C.R., Mirchandani, D., Scharton, D., Bilello, J.P., Ku, Z., An, Z., Kalveram, B., Freiberg, A.N., Menachery, V.D., Xie, X., Plante, K.S., Weaver, S.C., Shi, P.-Y., 2021. Spike mutation D614G alters SARS-CoV-2 fitness. Nature 592, 116–121. https://doi.org/10.1038/s41586-020-2895-3

Sailer, Z.R., Harms, M.J., 2017. Detecting High-Order Epistasis in Nonlinear Genotype-Phenotype Maps. Genetics 205, 1079–1088. https://doi.org/10.1534/genetics.116.195214

Schlub, T.E., Smyth, R.P., Grimm, A.J., Mak, J., Davenport, M.P., 2010. Accurately Measuring Recombination between Closely Related HIV-1 Genomes. PLOS Comput. Biol. 6, e1000766. https://doi.org/10.1371/journal.pcbi.1000766

Sharkey, M.E., Teo, I., Greenough, T., Sharova, N., Luzuriaga, K., Sullivan, J.L., Bucy, R.P., Kostrikis, L.G., Haase, A., Veryard, C., Davaro, R.E., Cheeseman, S.H., Daly, J.S., Bova, C., Ellison, R.T., Mady, B., Lai, K.K., Moyle, G., Nelson, M., Gazzard, B., Shaunak, S., Stevenson, M., 2000. Persistence of episomal HIV-1 infection intermediates in patients on highly active antiretroviral therapy. Nat. Med. 6, 76–81. https://doi.org/10.1038/71569

Starr, T.N., Czudnochowski, N., Liu, Z., Zatta, F., Park, Y.-J., Addetia, A., Pinto, D., Beltramello, M., Hernandez, P., Greaney, A.J., Marzi, R., Glass, W.G., Zhang, I., Dingens, A.S., Bowen, J.E., Tortorici, M.A., Walls, A.C., Wojcechowskyj, J.A., De Marco, A., Rosen, L.E., Zhou, J., Montiel-Ruiz, M., Kaiser, H., Dillen, J., Tucker, H., Bassi, J., Silacci-Fregni, C., Housley, M.P., di Iulio, J., Lombardo, G., Agostini, M., Sprugasci, N., Culap, K., Jaconi, S., Meury, M., Dellota, E., Abdelnabi, R., Foo, S.-Y.C., Cameroni, E., Stumpf, S., Croll, T.I., Nix, J.C., Havenar-Daughton, C., Piccoli, L., Benigni, F., Neyts, J., Telenti, A., Lempp, F. A., Pizzuto, M.S., Chodera, J.D., Hebner, C.M., Virgin, H.W., Whelan, S.P.J., Veesler, D., Corti, D., Bloom, J.D., Snell, G., (2021a). SARS-CoV-2 RBD antibodies that maximize breadth and resistance to escape. Nature 597, 97–102. https://doi.org/10.1038/s41586-021-03807-6

Starr, T.N., Greaney, A.J., Addetia, A., Hannon, W.W., Choudhary, M.C., Dingens, A.S., Li, J.Z., Bloom, J.D., (2021b). Prospective mapping of viral mutations that escape antibodies used to treat COVID-19. Science 371, 850–854. https://doi.org/10.1126/science.abf9302

Starr, T.N., Greaney, A.J., Hilton, S.K., Ellis, D., Crawford, K.H.D., Dingens, A.S., Navarro, M.J., Bowen, J.E., Tortorici, M.A., Walls, A.C., King, N.P., Veesler, D., Bloom, J.D., 2020. Deep Mutational Scanning of SARS-CoV-2 Receptor Binding Domain Reveals Constraints on Folding and ACE2 Binding. Cell 182, 1295–1310.e20. https://doi.org/10.1016/j.cell.2020.08.012

Starr, T.N., Greaney, A.J., Stewart, C.M., Walls, A.C., Hannon, W.W., Veesler, D., Bloom, J.D., 2022. Deep mutational scans for ACE2 binding, RBD expression, and antibody escape in the SARS-CoV-2 Omicron BA.1 and BA.2 receptor-binding domains. https://doi.org/10.1101/2022.09.20.508745

Tan, T.J.C., Mou, Z., Lei, R., Ouyang, W.O., Yuan, M., Song, G., Andrabi, R., Wilson, I.A., Kieffer, C., Dai, X., Matreyek, K.A., Wu, N.C., 2022. High-throughput identification of prefusion-stabilizing mutations in SARS-CoV-2 spike. https://doi.org/10.1101/2022.09.24.509341

Turakhia, Y., Maio, N.D., Thornlow, B., Gozashti, L., Lanfear, R., Walker, C.R., Hinrichs, A.S., Fernandes, J.D., Borges, R., Slodkowicz, G., Weilguny, L., Haussler, D., Goldman, N., Corbett-Detig, R., 2020. Stability of SARS-CoV-2 phylogenies. PLOS Genet. 16, e1009175. https://doi.org/10.1371/journal.pgen.1009175

Turakhia, Y., Thornlow, B., Hinrichs, A.S., De Maio, N., Gozashti, L., Lanfear, R., Haussler, D., Corbett-Detig, R., 2021. Ultrafast Sample placement on Existing tRees (UShER) enables real-time phylogenetics for the SARS-CoV-2 pandemic. Nat. Genet. 53, 809–816. https://doi.org/10.1038/s41588-021-00862-7

Van Maele, B., De Rijck, J., De Clercq, E., Debyser, Z., 2003. Impact of the Central Polypurine Tract on the Kinetics of Human Immunodeficiency Virus Type 1 Vector Transduction. J. Virol. 77, 4685–4694. https://doi.org/10.1128/JVI.77.8.4685-4694.2003

VanderPlas, J., Granger, B.E., Heer, J., Moritz, D., Wongsuphasawat, K., Satyanarayan, A., Lees, E., Timofeev, I., Welsh, B., Sievert, S., 2018. Altair: Interactive Statistical Visualizations for Python. J. Open Source Softw. 3, 1057. https://doi.org/10.21105/joss.01057

Wang, Q., Guo, Y., Iketani, S., Nair, M.S., Li, Z., Mohri, H., Wang, M., Yu, J., Bowen, A.D., Chang, J.Y., Shah, J.G., Nguyen, N., Chen, Z., Meyers, K., Yin, M.T., Sobieszczyk, M.E., Sheng, Z., Huang, Y., Liu, L., Ho, D.D., (2022a). Antibody evasion by SARS-CoV-2 Omicron subvariants BA.2.12.1, BA.4 and BA.5. Nature 608, 603–608. https://doi.org/10.1038/s41586-022-05053-w

Wang, Q., Iketani, S., Li, Z., Guo, Y., Yeh, A.Y., Liu, M., Yu, J., Sheng, Z., Huang, Y., Liu, L., Ho, D.D., (2022b). Antigenic characterization of the SARS-CoV-2 Omicron subvariant BA.2.75. Cell Host Microbe. https://doi.org/10.1016/j.chom.2022.09.002

Westendorf, K., Wang, L., Žentelis, S., Foster, D., Vaillancourt, P., Wiggin, M., Lovett, E., Lee, R. van der, Hendle, J., Pustilnik, A., Sauder, J.M., Kraft, L., Hwang, Y., Siegel, R.W., Chen, J., Heinz, B.A., Higgs, R.E., Kallewaard, N., Jepson, K., Goya, R., Smith, M.A., Collins, D.W., Pellacani, D., Xiang, P., Puyraimond, V. de, Ricicova, M., Devorkin, L., Pritchard, C., O’Neill, A., Dalal, K., Panwar, P., Dhupar, H., Garces, F.A., Cohen, C., Dye, J., Huie, K.E., Badger, C.V., Kobasa, D., Audet, J., Freitas, J.J., Hassanali, S., Hughes, I., Munoz, L., Palma, H.C., Ramamurthy, B., Cross, R.W., Geisbert, T.W., Menacherry, V., Lokugamage, K., Borisevich, V., Lanz, I., Anderson, L., Sipahimalani, P., Corbett, K.S., Yang, E.S., Zhang, Y., Shi, W., Zhou, T., Choe, M., Misasi, J., Kwong, P.D., Sullivan, N.J., Graham, B.S., Fernandez, T.L., Hansen, C.L., Falconer, E., Mascola, J.R., Jones, B.E., Barnhart, B.C., 2022. LY-CoV1404 (bebtelovimab) potently neutralizes SARS-CoV-2 variants. https://doi.org/10.1101/2021.04.30.442182

Yu, T.C., Thornton, Z.T., Hannon, W.W., DeWitt, W.S., Radford, C.E., Matsen, F.A., Bloom, J.D., 2022. A biophysical model of viral escape from polyclonal antibodies. https://doi.org/10.1101/2022.09.17.508366

Zhang, J., Cai, Y., Xiao, T., Lu, J., Peng, H., Sterling, S.M., Walsh, R.M., Rits-Volloch, S., Zhu, H., Woosley, A.N., Yang, W., Sliz, P., Chen, B., 2021. Structural impact on SARS-CoV-2 spike protein by D614G substitution. Science 372, 525–530. https://doi.org/10.1126/science.abf2303

Zhao, X., Zheng, S., Chen, D., Zheng, M., Li, X., Li, G., Lin, H., Chang, J., Zeng, H., Guo, J.-T., 2020. LY6E Restricts Entry of Human Coronaviruses, Including Currently Pandemic SARS-CoV-2. J. Virol. 94, e00562–20. https://doi.org/10.1128/JVI.00562-20

Zheng, M., Zhao, X., Zheng, S., Chen, D., Du, P., Li, X., Jiang, D., Guo, J.-T., Zeng, H., Lin, H., 2020. Bat SARS-Like WIV1 coronavirus uses the ACE2 of multiple animal species as receptor and evades IFITM3 restriction via TMPRSS2 activation of membrane fusion. Emerg. Microbes Infect. 9, 1567–1579. https://doi.org/10.1080/22221751.2020.1787797

Zhou, P., Song, G., He, W., Beutler, N., Tse, L.V., Martinez, D.R., Schäfer, A., Anzanello, F., Yong, P., Peng, L., Dueker, K., Musharrafieh, R., Callaghan, S., Capozzola, T., Yuan, M., Liu, H., Limbo, O., Parren, M., Garcia, E., Rawlings, S.A., Smith, D.M., Nemazee, D., Jardine, J.G., Wilson, I.A., Safonova, Y., Rogers, T.F., Baric, R.S., Gralinski, L.E., Burton, D.R., Andrabi, R., 2022. Broadly neutralizing anti-S2 antibodies protect against all three human betacoronaviruses that cause severe disease. https://doi.org/10.1101/2022.03.04.479488

Zufferey, R., Dull, T., Mandel, R.J., Bukovsky, A., Quiroz, D., Naldini, L., Trono, D., 1998. Self-Inactivating Lentivirus Vector for Safe and Efficient In Vivo Gene Delivery. J. Virol. 72, 9873–9880. https://doi.org/10.1128/JVI.72.12.9873-9880.1998

